# Scalable production of tissue-like vascularised liver organoids from human PSCs

**DOI:** 10.1101/2020.12.02.406835

**Authors:** Sean P Harrison, Richard Siller, Yoshiaki Tanaka, Yangfei Xiang, Benjamin Patterson, Henning Kempf, Espen Melum, Kathrine S Åsrud, Maria E Chollet, Elisabeth Andersen, Per Morten Sandset, Saphira Baumgarten, Flavio Bonanini, Dorota Kurek, Santosh Mathapati, Runar Almaas, Kulbhushan Sharma, Steven R Wilson, Frøydis S Skottvoll, Ida C Boger, Inger L Bogen, Tuula A Nyman, Jun J Wu, Ales Bezrouk, Dana Cizkova, Jaroslav Mokry, Robert Zweigerdt, In-Hyun Park, Gareth J Sullivan

## Abstract

A lack of physiological parity between 2D cell culture and *in vivo*, has paved the way towards more organotypic models. Organoids exist for a number of tissues, including the liver. However, current approaches to generate hepatic organoids suffer drawbacks, including a reliance on extracellular matrices (ECM), the requirement to pattern in 2D culture, costly growth factors and a lack of cellular diversity, structure and organisation. Current hepatic organoid models are generally simplistic, composed of hepatocytes or cholangiocytes, which renders them less physiologically relevant when compared to native tissue. Here we aim to address these drawbacks. To address this, we have developed an approach that does not require 2D patterning, is ECM independent combined with small molecules to mimic embryonic liver development that produces massive quantities of liver like organoids. Using single-cell RNA sequencing and immunofluorescence we demonstrate a liver-like cellular repertoire, a higher order cellular complexity, presenting with vascular luminal structures, innervation and a population of resident macrophage – the Kupffer cells. The organoids exhibit key liver functions including drug metabolism, serum protein production, coagulation factor production, bilirubin uptake and urea synthesis. The organoids can be transplanted and maintained in mice producing human albumin long term. The organoids exhibit a complex cellular repertoire reflective of the organ, have *de novo* vascularization and innervation, enhanced function and maturity. This is a pre-requisite for a myriad of applications from cellular therapy, tissue engineering, drug toxicity assessment, disease modeling, to basic developmental biology.

## INTRODUCTION

The liver is the largest endocrine organ in the body and is critical for maintaining homeostasis, serving as the primary site of xenobiotic metabolism, production of coagulation factors, and removal of ammonia as well as a multitude of other essential functions ^1^. In the setting of liver failure, no other curative treatment approach besides liver transplantation (LTX) exists. The gold standard for the evaluation of hepatic metabolism and drug toxicity, amongst other things, are primary human hepatocytes (PHHs). However these are limited in supply and rapidly lose function *in vitro*. This highlights the necessity to identify potential surrogates to fill this void. Human pluripotent stem cells (hPSCs) have the ability to self-organise into organotypic structures called organoids. However, organoid models do not fully recapitulate the cellular diversity and architectural features of the organ in question ^2^. The lack of such organotypic *in vitro* models has lead us to develop an approach that utilises a suspension culture system that leverages off the ability of hPSCs to self aggregate on seeding as single cells. In addition we have overcome the requirement to pattern the hPSCs in a 2D format to definitive endoderm (Ouchi et al., 2019) by directly patterning the 3D hPSC aggregates with the addition of small molecules, in an extracellular matrix (ECM) independent manner that is scalable to generate “mini-liver” organoids which contain both vasculature and innervation while recapitulating the complexity of cell types associated with the liver.

## RESULTS

### Differentiation of human pluripotent stem cells to liver organoids

This approach used a cocktail of small molecule mimetics in a developmentally relevant sequence to mimic *in vivo* liver development (Fig. 1A)^3–5^. On initiating organoid formation, we observed a rapid aggregation of the hPSCs post seeding, generating aggregates with an average diameter of 116 ±41 μm, while maintaining pluripotency markers (OCT4, SOX2 and NANOG) over 24 hours (Fig. 1B). To initiate the developmental programme, aggregates were challenged with a pulse of WNT signalling, via CHIR99021 (CHIR), which lead to an exit from pluripotency and a transition through primitive streak, resulting in aggregates of 128 ±57 μm (24 hours post CHIR treatment). The WNT signal was removed after 24 hours and by day 2 the aggregates exhibited markers of both mesoderm and definitive endoderm (DE); *T*, *GSC*, *FOXA2*, *SOX17*, *HHEX* and *CER1 etc* (Fig. 1C and D). During the initial 2 days of the differentiation we observed a shift in size of the organoids, resulting in mesendodermal aggregates with an average size of 159 ±46 μm (Fig. 1E). The mesendodermal aggregates were then subjected to hepatic specification until day 7, the resulting organoids were characterised for the presence of hepatic markers. We observed increased expression of *AFP*, *CEBPa*, *HNF4α* and *TTR* coinciding with decreased expression of DE markers such as *HHEX*, *SOX17* and *GATA4* (Fig. 1F). Expression of the T-box family protein *TBX3* was observed (Fig. 1F), which is a factor involved in hepatic endoderm delamination and subsequent invasion into the adjacent *septum transversum* mesenchyme (STM), resulting in the mixing of these two germ layers and the formation of the liver bud. Using immunostaining we revealed further characteristics of a mixed lineage liver bud stage. The outer epithelial layer of cells expressed ECAD (CDH1), FOXA2, and CK8, as well as being HNF4α and AFP positive, indicative of an early hepatic phenotype (Fig. 1G and Supplementary Fig. 1). These epithelial markers were localised to the outer surface and not the core of the organoids (Supplementary Fig. 1).

**Figure 1.**
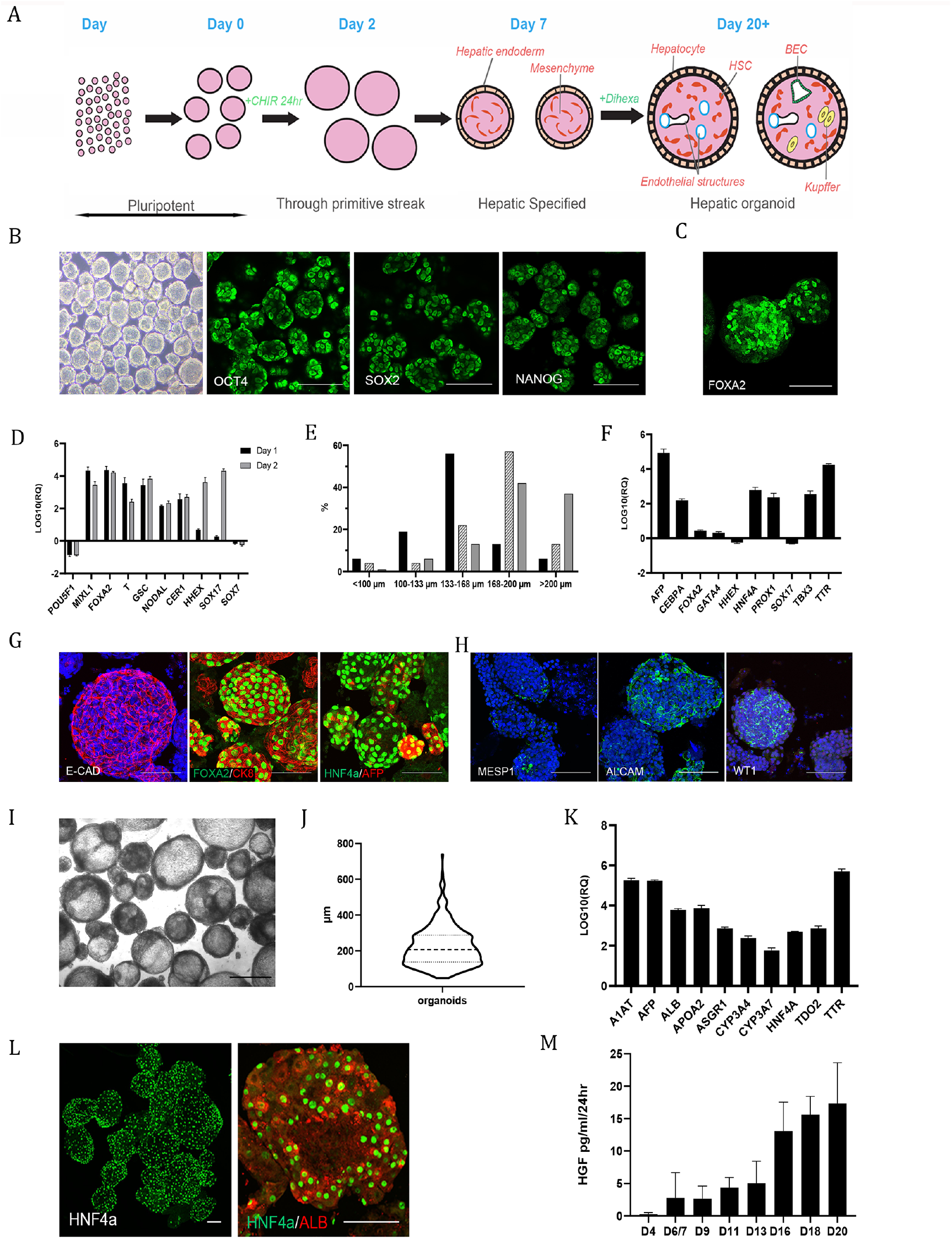
Differentiation of multicellular liver organoids from human PSCs, mimics stages of *in vivo* development. (A) Schematic overview of organoid differentiation from PSCs. (B) Representative images of Day 0 (D0) pluripotent spheroids, left panel brightfield. Remaining panels show whole-mount immunostaining of the pluripotency markers OCT4, SOX2 and NANOG. (C) Whole-mount immunostaining of the definitive endoderm (DE) marker FOXA2 at D2 of the differentiation. (D) RT-qPCR analysis of pluripotency and DE associated genes at D1 and D2 of the differentiation relative to D0 spheroids on a log10 scale. Results from three independent experiments are presented as mean ±SD. (E) Graph showing shift in size of organoids from D0 to D2. The size is expressed as a mean diameter (μm). The range from 100 to 200 μm was divided into three size groups. The percentages are based upon the amount in each category (represents a total number of 570 organoids). (F) RT-qPCR analysis of DE and early liver development genes at D7 of the differentiation relative to D2, on a log10 scale. Results from three independent experiments are presented as mean ±SD. (G) Whole-mount immunostaining of D7 organoids showing epithelial (ECAD) and early hepatocyte markers (FOXA2, CK8, HNF4α, AFP) on the outer surface of the organoids. (H) Whole-mount immunostaining of D7 organoids showing heterogeneous expression of mesoderm (MESP1) and mesenchymal (ALCAM, WT1) associated markers. (I) Brightfield image of D20 organoids scale bar is 500 μm. (J) Violin-plot of D20-D27 organoid diameter from three representative experiments, solid line represents the mean while dotted lines represent represent the quartiles. (K) RT-qPCR analysis of both early and later (developmentally) hepatocyte genes at D20 of the differentiation relative to D2, on a log10 scale. Results from three independent experiments are presented as mean ±SD. (L) Whole-mount immunostaining of D20 organoids showing expression of the hepatocyte markers HNF4α and Albumin on the outer surface of the organoids. (M) Demonstration of increasing secretion of HGF into the culture medium by the organoids throughout the differentiation as measured by ELISA, results from two independent experiments are presented as mean ±SD. All Scale bars are 100 μm unless stated otherwise.

We characterised the non-epithelial cores of day 7 organoids, where we observed a population of MESP1 positive cells, indicative of mesodermal STM ^6^(Fig. 1H). During development a mesodermal MESP1^+^ population gives rise to mesothelial/sub-mesothelial populations marked by ALCAM and Wilms Tumour (WT1), which in turn give rise to the hepatic stellate cells (HSCs) ^6, 7^. We also identified ALCAM and Wilms Tumour (WT1) positive populations within the mesenchyme of the organoids, these cells mark the putative mesothelial/sub-mesothelial populations (Fig. 1H). We then directed these early hepatic organoids to a mature liver-like stage over a 14-day period resulting in the formation of organotypic 3D structures (Figure 1I) with an average size of 250 μm (Fig. 1J). The liver organoids exhibited markers of hepatic maturation and tissue-like complexity. Analysis by RT-qPCR clearly demonstrated the expression of hepatic markers such as *ALB*, *A1AT*, *ASGPR1* as well as genes enriched in the liver such as *APOA2*, *TDO2* and *TTR* and those involved in xenobiotic metabolism cytochrome P450 (*CYP) 3A4* and *CYP3A7* (Fig. 1K). Immunofluorescence staining demonstrated that the hepatocytes were located on the surface of the organoids visualised *via* HNF4α and ALB staining (Fig. 1L). It is established that organogenesis is initiated when the epithelium interacts with an early mesenchymal population during liver development ^8^. This process is driven by a number of paracrine factors including the essential morphogen, hepatocyte growth factor (HGF) ^9^. We assessed the production of HGF during the course of the differentiation and noted secretion of HGF increased throughout the organoid differentiation (Fig. 1M). This gives further support to the developmental accuracy of the organoid differentiation as HGF is secreted early in development by the mesenchymal population and later by their derivative, e.g. the hepatic stellate cells (HSCs). This is shown *in vivo* to drive the expansion and maturation of the liver bud suggesting it fulfils a similar role here in our organoids ^10^.

### Single cell transcriptome analysis of iPSC derived liver organoids

To further dissect the cellular diversity within the liver organoids, we profiled the single-cell transcriptomes of a total of 21,412 cells by scRNAseq. A total of 22 clusters were detected and systematically assigned into liver cell types by unique markers, Gene Ontology (GO) functions and reference transcriptome profiles (Fig. 2A and Supplementary Fig. 2A-E) ^11–15^. We identified three hepatocyte-like clusters (HEP1-3) with substantial expression of glycerolipid or cholesterol metabolic genes ^16^. In mammalian liver, hepatocytes are hexagonally arranged into hepatic lobules that display a gradient microenvironment of oxygen, nutrients, hormones and secreted proteins from pericentral to periportal zones ^17^. To evaluate the spatial heterogeneity of hepatocytes within the liver organoid, we performed Gene Set Enrichment Analysis (GSEA) of gene signatures for human periportal hepatocytes to individual cells in the hepatocyte clusters (Fig. 2B) ^18^. The enrichment of periportal gene signatures was significantly different across the three hepatocyte clusters. In particular, the HEP3 cluster was closest to the periportal zone, while HEP1 was more distant suggesting that gene expression zonation of hepatocytes is present in the liver organoids. Next, the endothelial cell (EC)-like clusters were characterized by ECM genes and reference transcriptome annotation (Fig 2C and D) and divided into liver sinusoidal (LSEC) and macrovascular endothelial cells (MVEC) by the expression pattern of vascularization markers (Fig. 2C). To infer continuous EC heterogeneity, we ordered cells from EC clusters along with their gene expression patterns (Fig. 2D). Six co-expression modules that are transiently expressed with the EC ordering were involved in different cellular events. The modules biased to LSEC included genes related to oxidative reaction and anti-inflammatory response, which are characteristics of the liver sinusoid ^19,20^. In contrast, genes involved in smooth muscle and calcium transport were significantly enriched in MVEC-biased gene modules. Interestingly, in addition to the above liver cell types, we also identified clusters expressing genes for the development of peripheral nervous system (*SOX2*, *PAX6*, *STMN2*) (Fig. 2A,E and F and Supplementary Fig. 2B and E). This is not surprising as the human liver is highly innervated ^21^, it has also been demonstrated that neural crest cells appear in the epiblast prior to emergence of definitive ectoderm and mesoderm ^22, 23^.

**Figure 2.**
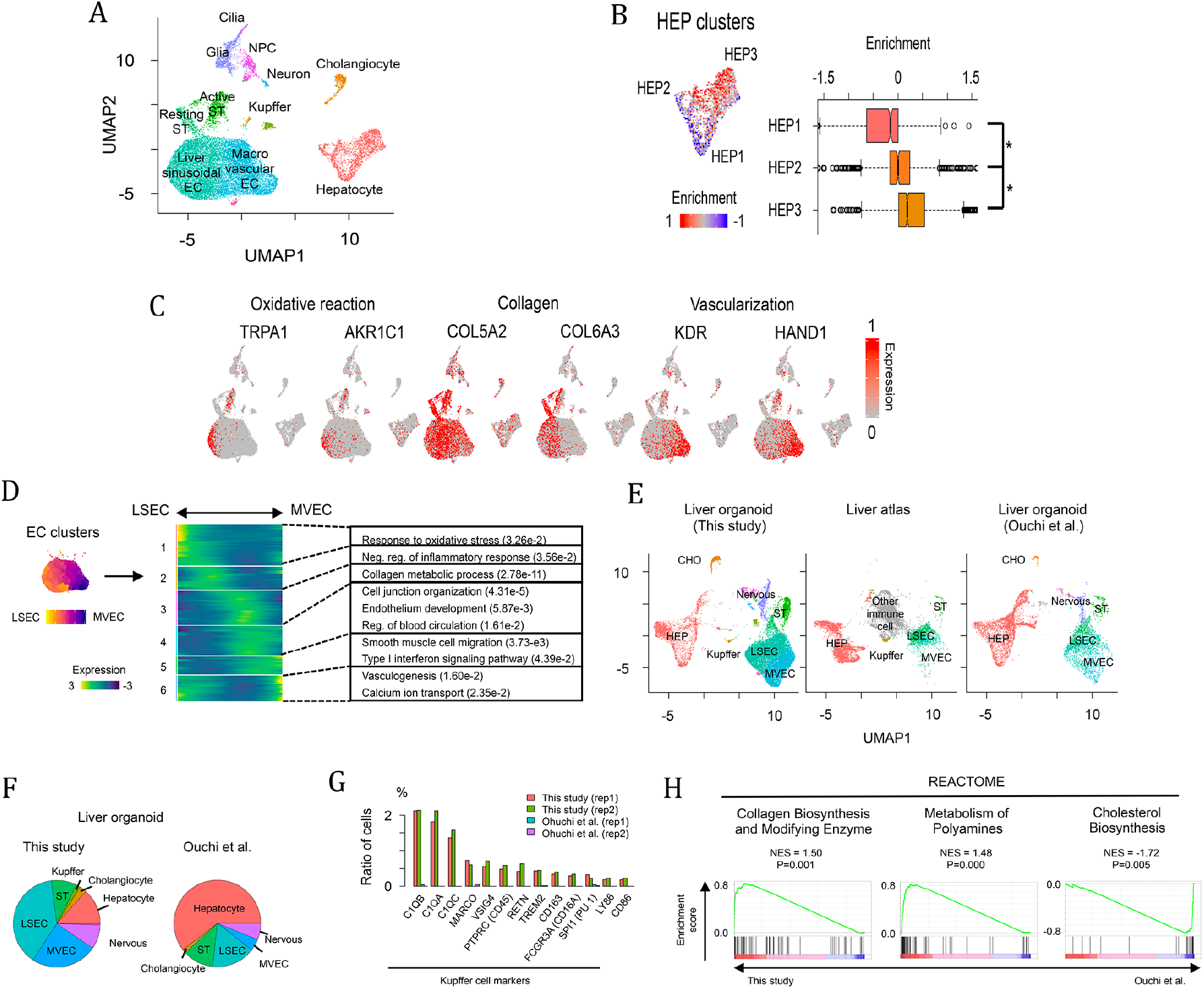
Single-cell RNA sequence analysis of liver organoids. (A) UMAP plot of single cells distinguished by cell types. (B) GSEA of gene signatures of periportal hepatocyte on hepatocyte clusters. The enrichment was compared across three hepatocyte clusters. * p<0.05 by two-sided student T test. (C) Expression pattern of endothelium-related genes in the liver organoid. Relative expression level is plotted from grey to red color. (D) Endothelial cell ordering from LSEC to MVEC. Heatmap represents expression pattern of genes, which are dependent on the EC ordering and categorized into six groups. Significant GO terms in each gene group are also shown. (E) UMAP plot of single cells derived from our and Ouchi *et al*. liver organoid^24^ and human liver^25^. (F) Pie charts representing cell composition in our and Ouchi *et al*. liver organoid. (G) Ratio of cells expressing Kupffer cell markers. (H) GSEA of pathway-related genes between our and Ouchi *et al*. liver organoid ^24^.

We next examined the consistencies and differences with primary liver and against another liver organoid protocol. Our scRNAseq data were compared against scRNAseq of FACS-sorted human adult liver cell populations and liver organoids derived from another protocol in the shared UMAP space (Fig. 2E and Supplementary Fig. 2F) ^24, 25^. The liver cell types (hepatocytes, HSCs, Kupffer and endothelial cells) in our liver organoid are close to cells derived from the human liver, indicating similar gene expression profiles of these cells between our organoid and primary human liver. The liver organoids from the two different protocols exhibited similar cell composition, but displayed apparent differences in the amount of non-endodermal cell types (Fig. 2F). For example, our protocol produces high number of endothelial cells, while more than half of cells are committed to hepatocytes in the Ouchi *et al*. organoids. In addition, peripheral nervous system-like cells were also detectable in the aforementioned study. Our comparative analysis revealed that the liver organoids displayed the generation and maturation of Kupffer cells (Fig. 2G) while not detectable in the Ouchi study. Cells expressing the cell surface marker (*MARCO*, *CD45*, *CD163*, *CD16A*, *LY86* and *CD86*), complement components (*C1QA*, *C1QB* and *C1QC*) and macrophage regulators (*VSIG4 TREM2* and *PU.1*) were clearly enriched in our protocol. For further comparison we analyzed the preferential pathways between the two protocols. GSEA for REACTOME database revealed that collagen and polyamine metabolic genes are significantly enriched in our protocol (Fig. 2H). In contrast, cholesterol biosynthetic genes were enriched in Ouchi and colleagues protocol, because of the higher number of hepatocytes in this protocol. Overall, our scRNAseq analysis indicates an improved organoid derivation.

### Proteomic analysis of iPSC derived liver organoids

Above we explored the composition of the organoids at the transcriptome level, next we subjected the organoids to global proteomic analysis. This revealed that 1,842 out of 2,461 detected proteins were expressed in both human primary liver (biopsies from human liver) and *in vitro* generated liver organoids. Using GO analysis these enriched proteins were involved in a variety of metabolic pathways and functions including blood coagulation (F2, SERPINC1, SERPING1, FGG, FGA etc.) and glutamine family amino acid metabolic processes (ARG1, ASS1, FAH, GLUL, GFPT1, GOT1; GOT2 etc.) (Fig. 3A). We identified approximately 200-400 proteins which were expressed differentially at the protein level (Fig. 3B) and found that those proteins related to blood cells (e.g. immune response and heme-binding) were significantly enriched in primary liver (Fig. 3B). In contrast, cell cycle and early developmental genes were enriched in the organoid.

**Figure 3.**
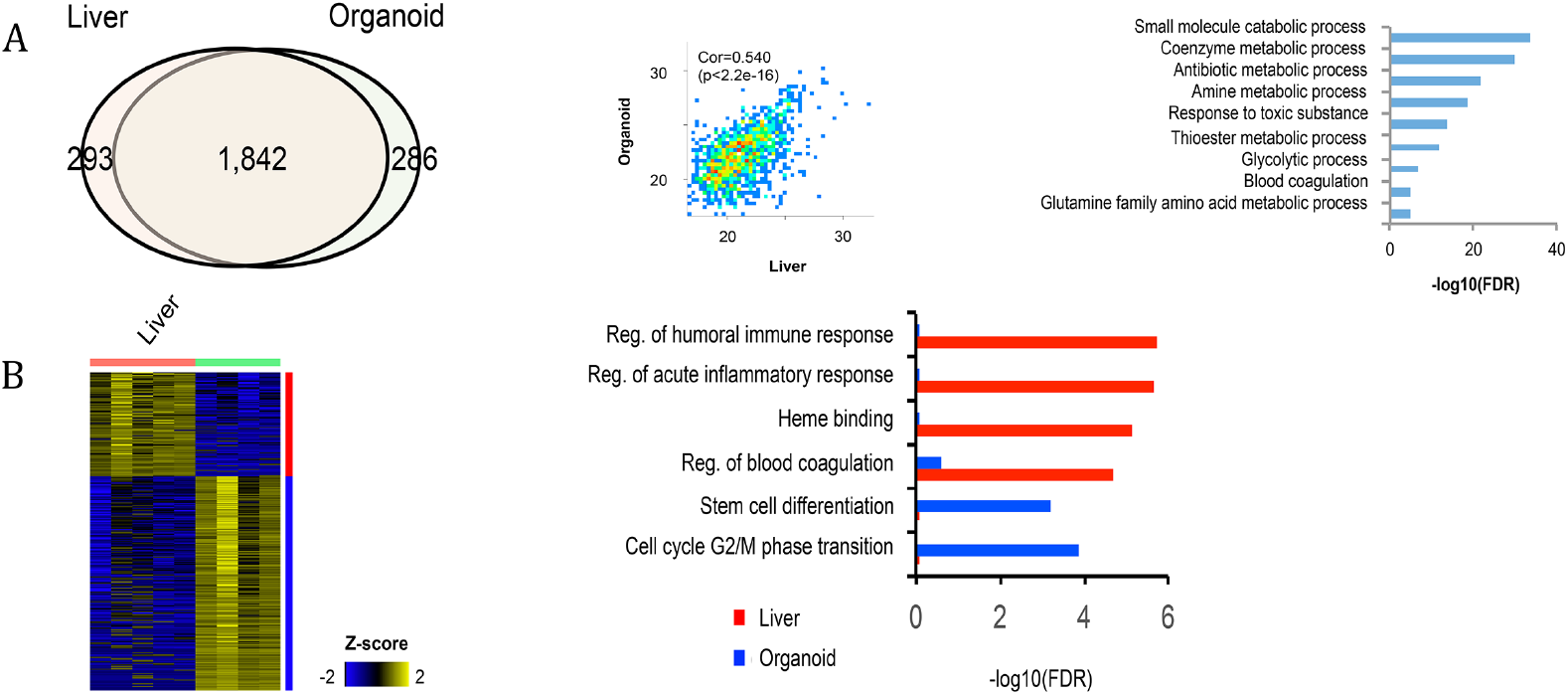
Proteomic analysis of liver organoids. (A) Venn diagram representing proteins shared between *in vivo* liver and *in vitro* organoid. The correlation is also shown with the scatterplot and the overrepresented GO terms of the shared proteins are shown by bar graph. (B) Differentially-expressed proteins and their overrepresented GO terms.

### iPSC derived liver organoids contain hepatic parenchymal cells

Next, immunofluorescence was performed to corroborate both scRNAseq and proteomic findings of liver-like cellular complexity. Organoids (day 20 to 30) were stained against a battery of typical hepatocyte markers. We verified the presence of hepatocytes by the cytosolic expression of GS and CPS1 along with nuclear expression of hepatic transcription factor HNF4α, which were located on the outer surface of the organoids (Fig. 4A). The enzymes, GS and CPS1, are involved in the urea cycle and in adult liver are zonally separated ^26^. The process of zonation occurs throughout postnatal development with GS being specifically absent from fetal liver hepatocytes ^27^. The expression of GS in our organoids is thus indicative of a developmental stage surpassing that of fetal liver. This is further reinforced by the expression of the xenobiotic metabolising enzyme CYP2A6 (Fig. 4A), a *bona fide* marker distinguishing adult from fetal hepatocytes ^28^ and the expression of ASGP1, a marker of maturity used to purify mature hepatocytes (Fig. 4A) ^29^. Another feature of hepatocytes is polarisation confirmed by expression of the tight junction protein ZO-1 and the apical export protein MRP2 (Fig. 4A). Polarisation was also assessed at the ultrastructural level, revealing features of primary liver tissue. This included epithelial cells lining luminal structures arranged in a layer and connected with tight junctions, a feature of the hematobiliary barrier. Along with a polarised epithelial cell morphology with the lumen facing surfaces presenting with numerous microvilli, while the abluminal surface facing the ECM was devoid of microvilli and attached to an underlying basal lamina (Fig. 4B). The other endoderm-derived cell type, the cholangiocyte, is derived from the hepatoblast and lines the biliary ducts that drain bile from the liver. We confirmed the presence of cholangiocytes, in cells surrounding lumen, through immunohistochemical staining of CK19 which is present in cholangiocytes and hepatoblasts but is lost in hepatocytes (Supplementary 3a) and CK7 which is expressed *in vivo* from 16-20 weeks post-conception (wpc) in humans (Fig. 4C) ^30^. Like the intestinal epithelium, the biliary epithelial cells produce mucins to form a mucus layer. We explored if we could detect the presence of mucus in our organoids. Using Alcian blue staining we observed epithelial cells lining the lumen stained blue indicating the presence of mucopolysaccharides i.e. mucin secreting cells (Fig. 4D). Interestingly cholangiocytes express neutral and acidic mucins from 23–40 wpc ^31^.

**Figure 4.**
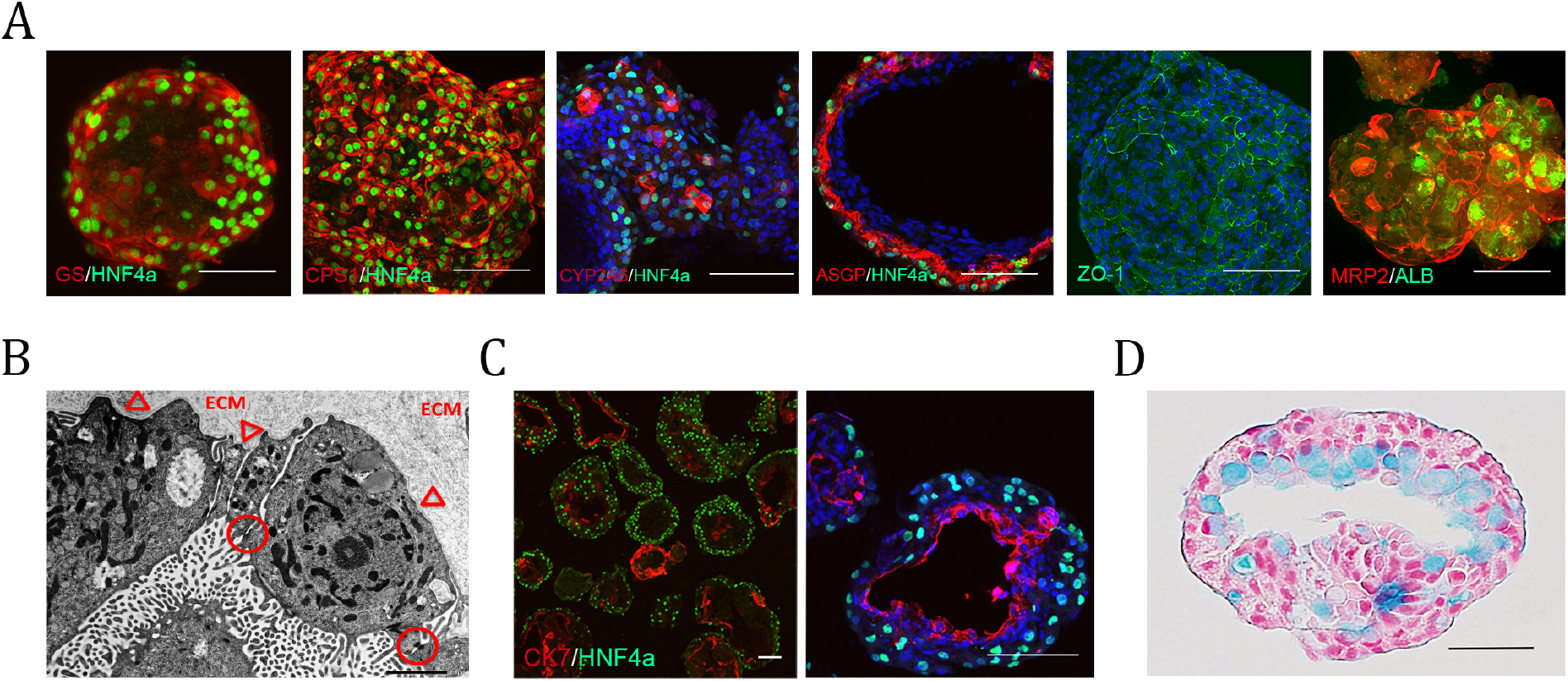
Liver organoids contain the parenchymal cell types of the liver. (A) Immunostaining showing expression of various maturity and polarity associated hepatocyte markers in the outer most layer of the organoids (whole-mount except for ASGP/HNF4α which are from 50 μm cryosections). (B) Electron micrograph revealing ultrastructural features associated with hepatocytes including epithelial cells lining a luminal structure arranged in one layer, connected with tight junctions (circles) indicative of polarization. The surface facing the lumen contains numerous microvilli whereas the abluminal surface facing the extracellular matrix remains smooth and attached to underlying basal lamina (arrowheads). Scale bar 2 μm. (C) Immunostaining of an organoid cryosection showing a later developmental cholangiocyte marker (CK7) positive population of cells separate to the hepatocyte population. These form smaller ring like structures as well as lining large luminal or cyst-like spaces within the organoids. (D) Paraffin embedded section of organoid stained with alcian blue and counterstained with nuclear red. The lumen of this organoid shows cells containing pale blue cytoplasm indicating the presence of mucopolysaccharides. Scale bar 50 μm.

### iPSC derived liver organoids are *de novo* vascularized and innervated

Along with the parenchymal cells the human liver is composed of a myriad of non-parenchymal cell types. We first investigated the endothelial populations identified by scRNAseq analysis. We used antibodies to delineate the different endothelial populations. First we investigated the macrovasculature and observed branched chains and lumen surrounding structures positive for CD31 (Fig. 5A and Supplementary Fig. 3b), we also observed small clusters of CD34^+^ endothelial structures suggesting continued neo-vascularization from approximately day 20 and beyond and not just the expansion of earlier endothelial structures (Fig. 5B). *In vivo*, the microvasculature in the liver bud is acquired from the endothelial cells of the STM, upon its invasion by hepatoblasts. These begin as CD34^+^/CD31^+^ continuous vasculature gradually acquiring liver sinusoidal endothelial cell (LSEC) specific features. In the adult liver LSECs exhibit distinct zonal markers ^32^, this is recreated in the organoids where the endothelial structures acquire increasing CD54^+^ expression in proximity to the hepatocyte layer, which suggests that endothelial cell specialisation is potentially a product of the niche (Fig. 5A and Supplementary Fig. 3c). We also investigated the distribution of LYVE1, another marker of LSECs associated with their scavenger function ^33^, this revealed luminal structures within the organoids, further confirming the heterogeneous and zonal nature of their vascularization (Fig. 5C).

**Figure 5.**
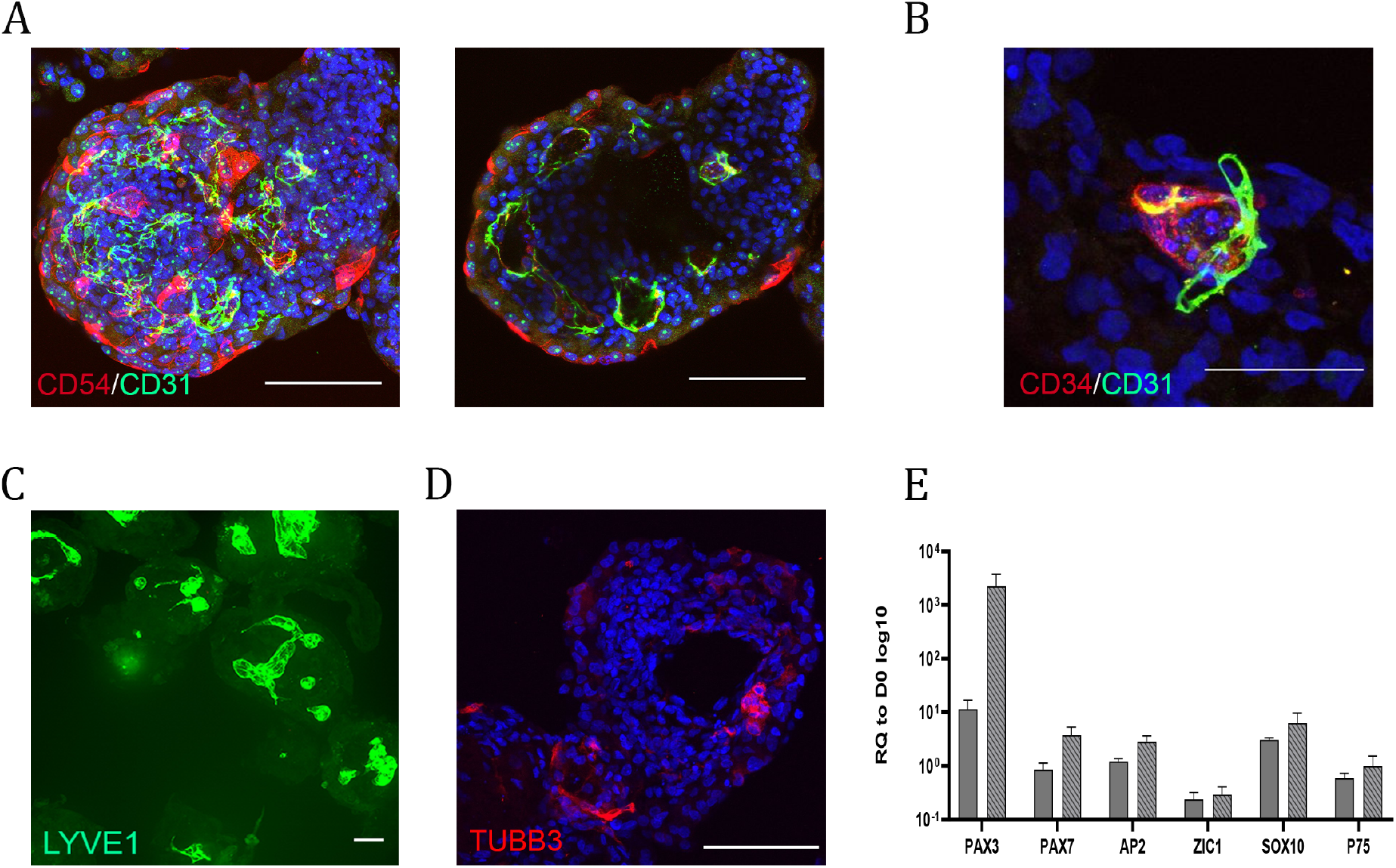
Liver organoids are *de novo* vascularised and innervated. (A) Max projection and cross section of whole-mount immunostained organoids showing overlapping populations expressing one or both of the endothelial markers CD54 and CD31. Cross-section reveals stronger CD54 expression towards the outer surface contrasted with stronger CD31 expression toward the centre of the organoid. Also visible are multiple conjoined luminal spaces bounded by the positive cells. (B), Immunostainng of a 50 μm cryosection showing adjacent and overlapping expression of CD34 and CD31 in a small structure indicative of neo-vascularization. (C) whole-mount staining of LYVE1 a sinusoidal endothelial marker showing positive cells demarking a diversity of lumen shapes and sizes. (D) Immunostaining for neuronal population in an organoid (TUBB3). (E) RT-qPCR analysis of neural crest stem cells markers, at D2 and D7 of the differentiation relative to D0 spheroids. Results from three independent experiments are presented as mean ±SD.

scRNAseq data identified a peripheral neuron population (Fig. 2A,E and F and Supplementary Fig. 2B and E). In order to corroborate these findings we performed immunohistochemistry against TUBB3 (Fig 5D), we detected TUBB3 positive neurons throughout the organoids. Interestingly the neural crest lineage arises from the pluripotent epiblast, prior to definitive germ layer formation ^22, 23^. We investigated early points in the differentiation (day 2 and 7) for the emergence of a neural crest like population. Using RT-qPCR we observed the neural plate border markers *PAX3, PAX7, ZIC1* and the neural crest specifiers *AP2* and *SOX10*. At day 7 also we observed the neural crest stem cell marker *p75* (Fig. 5E).

### iPSC derived liver organoids contain a resident macrophage and hepatic stellate population

scRNAseq data identified a discrete population of Kupffer cells, which are the resident macrophage population of the liver and are derived *in situ* during the hematopoetic phase of liver development *in vivo* ^34, 35^. We investigated the presence of Kupffer cells within the organoids using the marker CD68 ^36^, observing a CD68^+^ population exhibiting typical cytoplasmic granule staining (Fig. 6A). The endothelial and hematopoetic cells share a common precursor early in development called the hemangioblast. We speculated that the endothelial and Kupffer cell types potentially arise from a common mesodermal population at an early stage of differentiation. To that end we studied early time points in organoid differentiation using RT-qPCR for orchestrators of hematopoietic commitment and observed expression of *RUNX1* on day 2, which is essential for hematopoietic commitment. By day 7 we observed the expression of both *RUNX1 and GATA2* a player, along with *RUNX1,* in the early hemangioblast core circuit (Fig. 6B) ^37^. We speculate that part of this population will go on to form hematopoietic stem cells, but are not yet specified to a myeloid lineage, from which Kupffer cells would ultimately emerge. Together these data suggest that the mesoderm by virtue of undergoing differentiation adjacent to the endodermal derived cells, facilitate the same role as the mesoderm in liver development *in vivo*. The scRNAseq analysis identified a putative hepatic stellate cell (HSC) populations via canonical markers such as *BGN*, *CTGF*, *TPM2*, *SPARC*, *IGFBP*, *TAGLN*, *DCN*, *CCL2*, *COL1A1,* etc. *In vivo* HSCs originate from the undifferentiated mesenchyme ^7^. Above ALCAM^+^, MESP1^+^ and WT1^+^ populations were observed in the organoids (potentially a mesothelial/submesothelial equivalent) (Fig. 1h). We speculate that the WT1^+^ population potentially gives rise to a HSC population. We first assessed αSMA, revealing positive cells seen in close proximity to the hepatocyte population, marked by HNF4α (Fig. 6C and Supplementary Fig. 3d). On closer inspection both HSCs with a star-like morphology, that had long cytoplasmic processes with fine branches, and cells resembling myofibroblasts were observed suggesting quiescent and activated populations. This is corroborated by the scRNAseq data, where both activated and resting populations were detected in UMAP space (Fig. 2a). We also cannot rule out that the HSCs are fetal in nature, as undifferentiated fetal HSCs express αSMA ^38^. One function of HSCs is the production of ECMs including the laminins. Immunohistochemistry using a pan-laminin antibody along with αSMA revealed a close relationship between laminin and HSCs (Fig. 6C).

**Figure 6.**
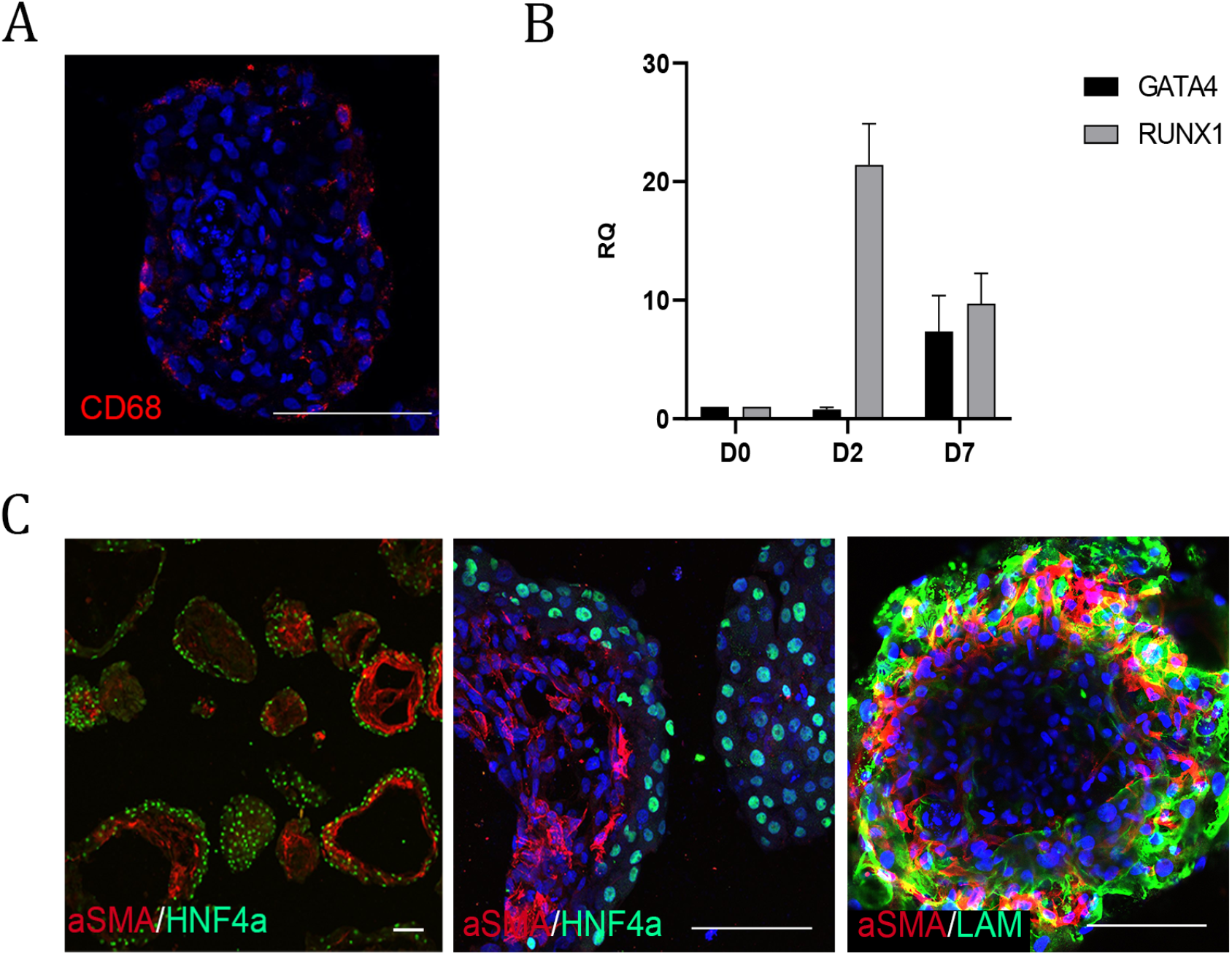
Liver organoids contain a resident macrophage and hepatic stellate population. (A) Immunostaining of 50 μm cryosection showing CD68 in a granular pattern in the cytoplasm of the Kupffer-like cells. (B) RT-qPCR analysis of hemangioblast (*RUNX1*) and hematopoietic (*GATA2*) associated genes involved in the development of macrophages, at D2 and D7 of the differentiation relative to D0 spheroids. Results from three independent experiments are presented as mean ±SD.(C) Immunostaining of cryosection showing expression of endodermal and stellate cell associated proteins HNF4a and αSMA (left and middle panel) and (right panel) whole-mount showing expression of mesenchymal and stellate cell associated proteins (αSMA, Laminin) beneath the hepatocyte layer of organoids. All Scale bars are 100 μm unless stated otherwise.

### iPSC derived liver organoids display liver like function

The organoids exhibit liver-like transcriptional and protein profiles, as well as a liver-like cellular repertoire. We next investigated the functional attributes of the organoids. The liver has a myriad of functions including the ability to metabolize drugs *via* the CYP450 enzymes. We assessed the basal and induced levels of CYP1A2 and CYP3A4, which play an important role in metabolism of a range of drugs including caffeine and acetaminophen ^39^. The organoids were benchmarked against a 2D protocol for generating hiPSC derived hepatocytes ^4, 5^. The basal and induced CYP activity in 2D cultures was similar to previous reports (day 20 of differentiation) ^4, 5^. The organoids presented with greatly elevated levels of activity for both basal and inducible metabolism at the equivalent time-point (Fig. 7A). These levels of activity (basal and induction) were maintained over a 40-day period in the organoids, while activity rapidly declined to almost undetectable levels in 2D culture (Fig. 7A). We then assessed long-term activity, observing maintenance of basal and inducible activity for 80 days (Fig. 7B). The signal started to decline from day 50 for CYP3A4 and day 60 for CYP1A2; we speculate that this may be a feature of suboptimal culture conditions for long-term maintenance, possibly due to changing mass transfer conditions within the organoids (Fig. 7B). The field of non-CYP450-mediated metabolism has attracted increasing attention as an important player in absorption, distribution, metabolism, and excretion ^40^. We investigated liver carboxyl esterases (CES), which are hydrolytic enzymes involved in the metabolism of endogenous esters, ester-containing drugs, pro-drugs and environmental toxicants ^41^. The CES enzymes also metabolize a wide range of xenobiotic substrates including heroin, which is metabolized by sequential deacetylation (phase I reaction) to 6-monoacetylmorphine (6-MAM) and morphine. Morphine is then glucuronidated, by a phase II reaction via UDP-glucuronosyltransferase (UGT) to morphine-3-glucuronide (M3G) (Fig. 7C). We tested if organoids supported heroin metabolism by exposure to 10 μM heroin and quantification by UPLC-MS/MS, whereupon we observed phase I metabolism (CES) to morphine, followed by phase II metabolism (UGT) to morphine glucuronides (Fig. 7C).

**Figure 7.**
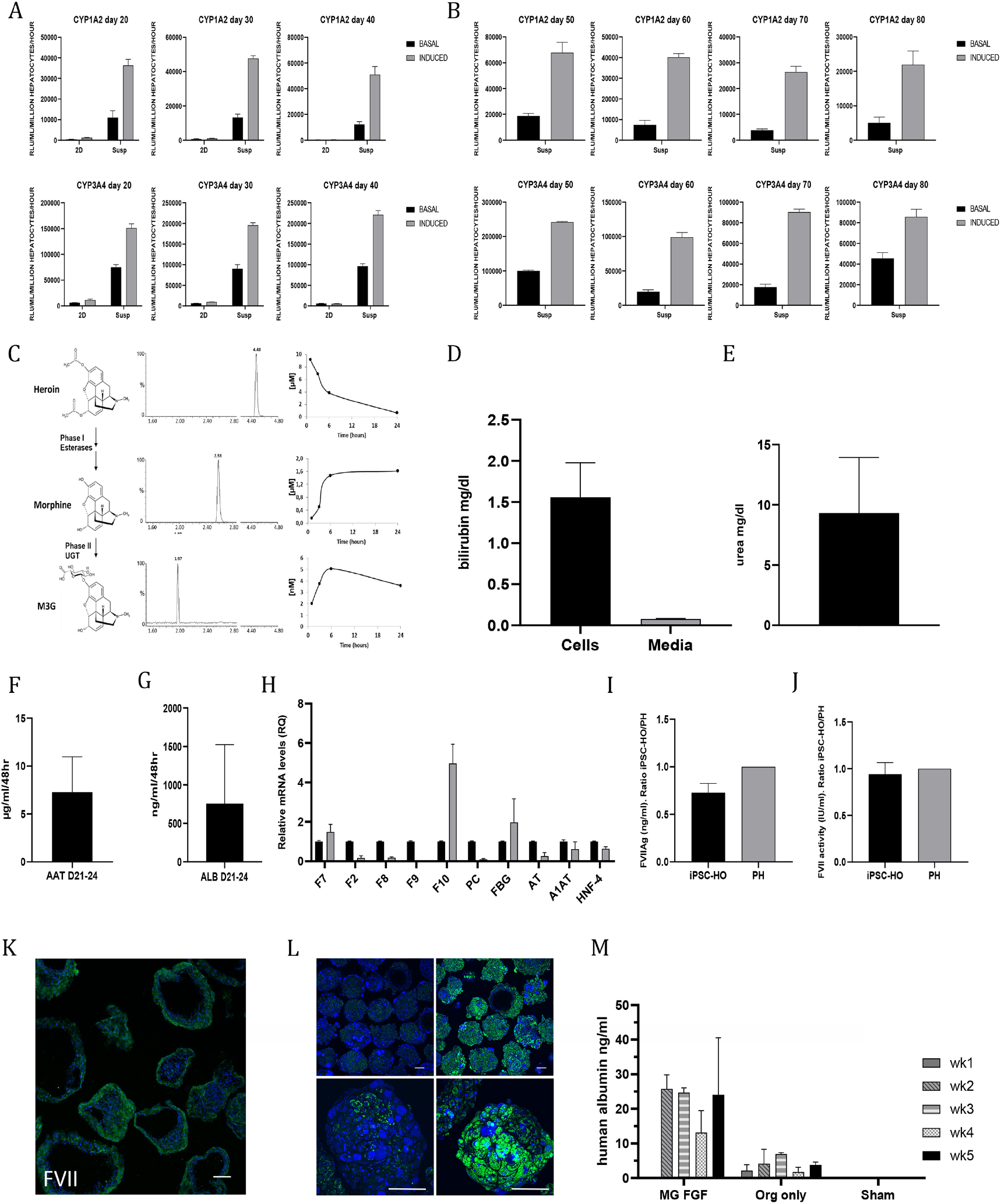
Assessment of liver organoid function, maintenance and transplantability. (A) Assay demonstrating inducibility and increasing activity of CYP1A2 (top row) and CYP3A4 (bottom row) drug metabolizing enzymes at D20, D30 and D40 in the hESC line H1 organoids. To induce CYP1A2 the cells were pretreated with omeprazole and for CYP3A4 rifampicin. Graphs show comparison of suspension cultures and 2D differentiation. Results from three independent experiments are presented as mean ±SD. (B) Assay demonstrating long-term maintenance of activity and inducibility of CYP1A2 (top row) and CYP3A4 (bottom row) proteins in suspension up to D80. Results from three independent experiments are presented as mean ±SD. (C) D20 Organoids show Phase I and Phase II metabolism of heroin dosed at 10 μM. Left panel represents the metabolic pathway of heroin. Heroin is metabolized by sequential deacetylation (phase I reaction)to 6-monoacetylmorphine and morphine by esterase enzymes. Morphine is further glucuronidated (phase II reaction) by UDP-glucuronosyltransferases (UGTs) to morphine-3-glucuronide (M6G) and morphine-6-glucuronide (M3G). Center panel; Ion chromatograms of an extracted organoid sample analysed by LC-MSMS. Right panel; Metabolism of heroin (10 μM) in organoids. (D) Demonstrating bilirubin (2 μM) uptake over 48 hours by D20 organoids after addition to the culture medium (n=3, mean ±SD). (E) Demonstrating the production and secretion of urea into the culture medium after 48 hours in D20 organoids (n=3, mean ±SD, no urea was detectable in cell-free medium). (F and G) Demonstrating the production and secretion of A1AT and albumin into the culture medium by D21-24 organoids over 48 hours as measured by ELISA (n=3, mean ±SD). (H) Levels of coagulation factors and inhibitors in iPSC derived organoids (gray bars) and primary human hepatocytes (black bars). mRNA levels of coagulation factors II, VII, VIII, IX, X, fibrinogen (*F*2, *F*7, *F*8, *F*9, *F*10, FBG), coagulation inhibitors protein C and antithrombin (PC, AT) and hepatic markers alpha-1 antitrypsin and hepatocyte nuclear factor 4 alpha (A1AT, HNF4α) were determined using quantitative RT-qPCR with 18S as endogenous control. The results are presented as mean of the fold-change expression of the respective gene. Results from three independent experiments are presented as mean ±SD. (I) Demonstrating the production and secretion of FVII protein in culture medium of iPSC derived organoids (iPSC-HO) and primary human hepatocytes (PH) was determined using ELISA. The total concentration of FVII was adjusted to 1×10^6^ cells and the results are expressed as the ratio iPSC-HO/PH. Results from three independent experiments are presented as mean ±SEM. (J) Demonstrating FVII activity (IU/ml), culture medium from iPSC-HO and PH was determined using a FVII chromogenic assay. Results were adjusted to 1×10^6^ cells and expressed as the ratio iPSC-HO/PH. Results from three independent experiments are presented as mean ±SEM. (K) Immunostaining of 50 μm cryosection showing localization of FVII to the outer hepatocyte layer of the organoids. (L) Whole-mount live imaging of Oleic acid treated (right column) and untreated (left column) organoids showing accumulation of non-polar fats after treatment (BODIPY in green). Oleic acid was used at 300 μM for 5 days. (M) Transplanted organoids can be maintained in mice. Human albumin was consistently detected over a 5 week period in mouse blood samples. D8 organoids were transplanted with either Matrigel / FGF2 supplementation or Matrigel / FGF2 free (mean ±SD; MG+FGF n=3, ORG n=2, sham n=4). All Scale bars are 100 μm unless stated otherwise.

The liver is also responsible for the clearance of bilirubin, the end product of heme catabolism, specifically this uptake is carried out by the hepatocytes ^42^. We investigated bilirubin uptake *via* a total bilirubin assay where we demonstrated the effective uptake of unconjugated bilirubin by organoids over a period 48 hours (Fig. 7D). This is not surprising as scRNAseq data shows the expression of the organic anion transporters in the hepatocyte population. Another essential liver function explored was hepatic urea synthesis, which is required for the removal of excess nitrogen. Above we show the expression of CPS1 and GS enzymes, which are involved in the urea cycle (Fig. 4A). The organoids produced and secreted urea into the medium at an average concentration of 9.3 mg/dl, while undetectable in the control (Fig. 7E), these values are within the range observed in blood of an adult (7-20 mg/dl) ^43^.

An important feature of the liver is the production of a plethora of serum proteins, including major plasma proteins, apolipoproteins, coagulation factors and hormones etc. We have demonstrated the production of HGF from our organoids (Fig. 1M); on inspecting the proteomic datasets numerous serum proteins were identified, including the apolipoproteins (APOA1, APOA4, APOC3 and APOD), hormones (the IGFs) and serine protease inhibitors (SERPING1 and SERPINA1/ alpha-1-antitrypsin (A1AT)) etc. Next, we investigated the production and secretion of albumin and A1AT, previously shown to be transcribed (Fig. 1K) and albumin expressed in hepatocytes (Fig. 1L). Both these proteins are secreted from hepatocytes into the circulation *in vivo* and their secretion into the culture medium was also verified (Fig. 7F and G. The liver also produces and secretes coagulation factors and inhibitors to maintain balanced hemostasis. We investigated if we could detect the expression of the vitamin K-dependent coagulation factors and inhibitors as well as a number of other coagulation factors. The expression of vitamin K-dependent coagulation factors (with the exception of *F9* which exhibited very low expression), inhibitors, *F8* (expressed in endothelial cells such as LSECs), *FBG* and *anti-thrombin* (*AT*) were all observed (Fig. 7H). Interestingly the mRNA levels of *F10* showed elevated expression compared to the other coagulation factors. We speculate that the elevated levels maybe a feature of the activated HSC population which (indicative of fibrosis) leads to elevated F10 ^44^ and worthy of further investigation.

We then investigated protein synthesis and secretion of the vitamin K-dependent coagulation factors with the highest mRNA levels (*F7* and *F10*) by ELISA and the intracellular levels of *F2* by western blot. We observed both the production and secretion of F7 and F10 into the cell medium (Fig. 7I and Supplementary Fig. 3e) and F2 in hepatic organoids lysates (Supplementary Fig. 3f). F7 has a key role in the initiation of blood coagulation so F7 activity was assessed, observing levels comparable to primary hepatocytes (Fig. 7J. Finally immunofluorescence demonstrated that F7 localized to the outer epithelial cells, corresponding to the hepatocytes (Fig. 7K).

A study from Ouchi *et al.* (2019) demonstrated the accumulation of lipids in a liver organoid model, which displayed a steatohepatitis-like phenotype ^25^. We tested whether accumulation of lipids could be established in our organoids. Using the neutral lipid dye Bodipy we established the steady-state level of lipids in untreated organoids, which was low (Fig. 7L). We next treated organoids with a free fatty acid, which lead to massive lipid accumulation, in enlarged droplets (Fig. 7L).

Our ECM independent system combined with small molecules capable of producing 300-500 organoids per ml of culture media, where we routinely produce 10’s to 100’s of thousands of organoids (see Supplementary Fig. 4a). Importantly, the cost of production has been reduced by nearly 3 orders compared to conventional 2D approaches (Supplementary Fig. 4b). With access to such numbers of organoids we assessed if our liver organoids could be transplanted and maintained in the kidney capsule of mice. The organoids were introduced into the kidney capsule with or without ECM. Using a human specific albumin assay as readout, we assessed the blood of mice over a 5 week period. No albumin was detectable in the first 96 hours, however we observed secretion of human albumin into the bloodstream of the recipient mice from weeks 1 to 2 post transplantation. This was maintained until the mice were sacrificed at week 5 post-transplantation, while no human albumin was detectable in the sham mice (Fig. 7M).

## DISCUSSION

The production of scalable liver-like tissue that exhibits functionally long lived human liver characteristics has remained elusive. Here we describe an approach that can generate massive amounts of functionally mature hPSC derived liver-like organoids. The described protocol is straightforward, efficient, reproducible and organoids can be produced in just 20 days. The organoid differentiation protocol follows a liver-like developmental route, producing organoids that contain a liver-like cellular repertoire including the parenchymal hepatocytes and cholangiocytes. They also contain non-parenchymal cells, hepatic stellate, Kupffer and endothelial populations. Interestingly, further analysis by scRNAseq indicated the presence of both hepatocyte and endothelial zonation. Proteomic analysis also clearly demonstrated a liver like phenotype. scRNAseq data was further confirmed using immunostaining, where we observed the aforementioned cell types. Interestingly, we observed parenchymal cell polarity, which was further supported by electron microscopy. We also confirmed the presence of the non-parenchymal cell types including the stellate and Kupffer cells. Remarkably we also observed *de novo* vascularization and innervation within our organoids. Finally the organoids present with liver like functional features, which included the production of serum proteins and the coagulation factors. They supported ureagenesis and bilirubin uptake. They are proficient in drug metabolism exhibiting long-term activity, 80 days, with respect to CYP metabolism. They also exhibited non-CYP mediated metabolism using heroin as an exemplar. The organoids have the ability to accumulate fatty acids, presenting with a steatotic like phenotype, potentially providing a model for NAFLD. Finally the organoids can be successfully transplanted into mice where they stably produce human albumin.

Organoids can provide both a unique and powerful model to interrogate disease, understand regeneration and potentially provide the building blocks for bridging therapies for patients waiting for an organ or ultimately provide a replacement organ. A key limitation is scaling, for example in children 10^9^ hepatocytes are required to correct specific metabolic liver function ^45^, however researchers are currently producing organoids in the 10’s to 100’s in combination with Matrigel (ECM) and recombinant growth factors, making scaling both a financial and technical bottleneck. Our approach currently produces around 500 organoids per ml of medium used (Supplementary Fig. 4a) in Erlenmeyer flasks. Therefore based on our approach we would require culture volumes of 1-3 litres to achieve these numbers. It is also compatible with the controlled production in stirred tank bioreactors as recently demonstrated for cardiac and hematopoietic lineages ^46, 47^ and will provide the field with a game-changing resource to allow the development of clinical as well as screening platforms where the requirement will be in the millions of organoids rather than todays laboratory scale.

More work is required to investigate the potential of these scalable organoids in the context of transplantation for bridging therapies but will potentially provide an important treatment modality for non-reversible acute liver failure ^48^. Timing of the transplantation is critical for acute liver failure and many patients receive a graft from a marginal donor instead of waiting for a better offer, which is associated with worse post-transplant outcome ^49, 50^, or succumb to disease without transplantation. Therefore approaches that can improve the condition and prolong the survival of patients with acute liver failure waiting for a liver transplant would increase the number of potential organ offers for a cadaveric graft, reducing the use of marginal donors and even allow transplants to be performed in patients currently not getting a graft offer. We envisage these organoids will provide a powerful tool to address developmental biology in the dish, where we could utilise lineage tracing to investigate the emergence of HSCs and the tissue-resident hematopoietic cells i.e. Kupffer population. Organoid production is scalable at a cost-effective level, but will require standardization in order to provide a platform for drug screening, toxicology, disease modeling, to act as the building blocks to produce liver micro-tissues and potentially scaling to large tissue units.

## MATERIALS AND METHODS

### hPSC Culture

The human pluripotent stem cell lines utilised in this study were as follows: the human embryonic stem cell line H1 (WiCell) and the previously described hiPSC lines AG27 (reprogrammed using retrovirus from AG05836B fibroblasts, obtained from Coriell Cell Repositories), and Detroit RA (reprogrammed using Sendai virus from Detroit 551 fibroblasts, obtained from ATCC) ^4, 5, 51, 52^. hPSCs were maintained under feeder free conditions on Geltrex (Life Technologies) coated tissue culture plates using Essential 8 medium made in house as described previously ^52^.

### Culture of primary human hepatocytes

Human plateable hepatocytes (primary hepatocytes (PH)) were purchased from Thermo Fisher Scientific and cultured in Williams’ Medium E (1x, no phenol red) (Thermo Fisher Scientific) following the manufacturer’s instructions.

### 2D hepatocyte-like-cell differentiation

Cells were differentiated to hepatocyte-like-cells (HLCs) in 2D as described previously ^4, 5, 52^. Briefly, cells were initially seeded onto Geltrex coated tissue culture plates as single cells after incubation with Accutase (Life Technologies). Optimal seeding density was previously established empirically as described in Mathapati et al., 2016^4^. Differentiation to HLCs was through a 3 stage process consisting of differentiation to definitive endoderm (DE—Phase I), hepatic specification to hepatoblasts / hepatic endoderm (HE—Phase II), and finally maturation to hepatocyte like cells (HLCs—Phase III). In order to ensure high-quality 2D differentiation we utilised our recently reported method for lifting and replating hepatic endoderm cells ^52^. For a detailed protocol of the differentiation process, please see Mathapati et al., 2016 and Siller et al., 2017^4, 52^.

### Suspension culture and differentiation to liver organoids

To differentiate hPSCs to organoids the cells were harvested by incubating with Accutase (Life Technologies) for 10 minutes at 37℃ until all cells had detached. The cells were pelleted by centrifugation at 300 × g for 5 minutes at room temperature. After counting, the cells were seeded into 125 ml or 500 ml cell culture Erlenmeyer flasks (Corning) at a bulk cell density of 3.5 - 4 ml of media/million cells, (see ^53^) in Essential 8 supplemented with 10 μM Y-27632 (BOC Sciences). The cells were allowed to self-organise into aggregates for 24 hours on an orbital shaker at 70 RPM in a humidified 37℃, 5% CO_2_ incubator. The conditions for suspension culture (orbital shaker at 70 RPM in a humidified 37℃, 5% CO_2_ incubator) were utilised for all further steps of the differentiation. After aggregate formation, differentiation was commenced to drive pluripotent hPSC aggregates to primitive streak / mesendoderm (day 1) and further patterned towards definitive endoderm (day 2) (See Fig. 1a for a schematic overview of the differentiation). To initiate differentiation, the hPSC aggregates were collected from the flask and transferred to a 50 ml conical tube and pelleted by centrifugation for 5 minutes at 300 × g at room temperature. After removal of the supernatant, the aggregates were resuspended in 3.5 ml/million cells of differentiation medium comprised of RPMI 1640 (Life Technologies) supplemented with B-27 either with or without Insulin (RPMI/B-27 +/−) (Life Technologies) depending on the cell line and 3 or 4 μM CHIR99021 (BOC Sciences). Optimal conditions need to be established for each line based on our previously established protocol ^4, 5, 51, 52^. The aggregates were then transferred back to the Erlenmeyer flask and incubated for another 24 hours. After 24 hours, the aggregates collected as previously described and the cell pellet was gently resuspended in the same volume of RPMI/B-27 +/−, without any small molecules, and transferred back to the Erlenmeyer flask. The aggregates were incubated for a further 24 hours. On day 2 of the differentiation the cells were directed towards hepatic endoderm (day 7). The aggregates were collected as above and resuspended at 3.5 ml/million cells in Knockout DMEM (Life Technologies), 20% (vol / vol) Knockout Serum Replacement (Life Technologies), 1% dimethyl sulfoxide (Sigma-Aldrich), non-essential amino acids (NEAA—Life Technologies), 2-mercaptoethanol (Life Technologies) and Glutamax (Life Technologies) and incubated for 5 days, with medium changes every 48 hours. On day 7, the resulting organoids were switched to medium for maturation to liver organoids. The medium comprised of Lebovitz L-15 base medium supplemented with 8.3% foetal bovine serum (FBS-Biowest), 8.3% Tryptose Phosphate Broth (Sigma-Aldrich), Hydrocortisone (Sigma-Aldrich), Ascorbic Acid (Sigma-Aldrich), Glutamax (Life Technologies), 100 nM Dexamethasone (Sigma-Aldrich), and 100 nM *N*-hexanoic-Tyr, Ile-6 aminohexanoic amide (dihexa) (Active Peptide). The organoids were cultured from day 7 to day 20, with a medium exchange every 48 hours. Organoids were collected and analysed or maintained in long-term culture as indicated.

### Fixation of organoids

For TEM the organoids were fixed in 1% glutaraldehyde / 1% paraformaldehyde (PFA) in 0.12 M phosphate buffer and 0.02 mM CaCl_2_ (pH 7.2 – 7.5; Sigma) for 4 hours at room temperature, washed in 8 % glucose (in 0.12 M phosphate buffer and 0.02 mM CaCl_2_, pH 7.2 – 7.5) and post-fixed in 2% OsO_4_ (in 0.12 M phosphate buffer and 0.02 mM CaCl_2_, pH 7.4; Sigma) for 90 minutes at room temperature. For histology and immunohistochemical detection the organoids were briefly rinsed in 0.1 M Sörensen buffer (pH 7.4) and immersed in 3% PFA and 0.05% glutaraldehyde in 0.1 M Sörensen buffer (pH 7.4) for 2 hours at room temperature followed by 30 minutes at 4°C. After a thorough washing in 0.1 M Sörensen buffer (pH 7.4), the organoids were dehydrated and embedded in paraffin. 6 μm thick serial sections were cut from paraffin blocks using a microtome and every tenth slide was stained with hematoxylin-eosin for histological examination.

Immunohistochemical detection of cytokeratin 18 and cytokeratin 19 was performed by indirect two-step method in paraffin-embedded sections. After deparaffinization and rehydration of sections, antigen retrieval was performed in HistoStation (Milestone, Sorisole, Italy). Endogenous peroxidase was blocked in 5% H_2_O_2_ (3 × 10 minutes) and then, sections were incubated with primary mouse anti-cytokeratin 18, clone DC10 (DAKO, Glostrup, Denmark; 1:25) or primary mouse anti-cytokeratin 19, clone BA17 (DAKO, Glostrup, Denmark; 1:50) antibody for 1 hour at room temperature. After washing in PBS, the sections were exposed to anti-mouse DAKO EnVision+ System-HRP Labeled Polymer (DAKO, Glostrup, Denmark) for 35 minutes at room temperature. Then the reaction was developed with 3,3-diaminobenzidine tetrahydrochloride (Sigma-Aldrich). Sections were dehydrated, counterstained with hematoxylin and mounted in DPX (Sigma-Aldrich). Tissue sections were examined in Olympus BX51 microscope equipped with DP71 camera.

For whole-mount immunofluorescence and confocal imaging organoids were fixed in 4% PFA for 60 minutes, then pelleted (300 × g for 5 minutes, low deceleration) and washed in PBS for 20 minutes, repeated three times. Fixed samples were stored in PBS at 4°C until required. 10 μl of sphere sediment was pipetted onto a coverslip and briefly air-dried followed by centrifugation at 1000 × g for 5 minutes. The fixed organoids were blocked for 1 hour in PBS supplemented with 0.1% Triton X-100 (Sigma Aldrich) × 100 (PBS- Triton X-100) and 10% goat serum (Life Technologies). The primary antibody was then diluted at the appropriate dilution in PBS- Triton X-100 containing 1% goat serum and incubated overnight at 4°C. The primary antibody was removed and the sample washed for 3 × 20 minutes in PBS-Triton X-100. All Alexa-Fluor secondary antibodies (Life Technologies) were diluted 1:1500 with PBS-Triton X-100 and added to the samples for 4 hours at 4°C in the dark and subsequently washed 3 times for 20 minutes in PBS-Triton X-100. To dual label the samples, the above steps (block, primary, secondary) were repeated. Nuclei were counterstained with DRAQ5 at 1:1500 in PBS, for a minimum of 15 minutes before imaging.

### Microscopy

Phase-contrast images were obtained using an Axio Primovert upright light microscope (Zeiss). Images were captured using Zen Software (Zeiss). All scale bars represent 100 μm, unless otherwise stated in the figure legends. Confocal images were obtained using an Olympus FV1000 confocal microscope with submersion lenses, images were captured using Olympus Fluoview and compiled using Image J software. Scale bars represent 100 μm unless otherwise stated in the figure legends.

### Cryosectioning

PFA fixed organoids were transferred to a cryomold and embedded in OCT compound (Thermo Fisher Scientific) and cooled to −80°C on a bath of isopropanol on dry ice. The cryo-embedded organoids were then sectioned at 50 μm on a cryotome and transferred to slides. Slides were stored at −80°C until immunostained and imaged as described above.

### Transmission electron microscopy

The fixed organoids (described above) were rinsed and incubated overnight in 10% sucrose (in water) at 4°C, the organoids were dehydrated in graded alcohols (50%, 75%, 96%, 100%), cleared in propylene oxide and embedded in a mixture of Epon 812 and Durcupan (Sigma; polymerization for 3 days at 60°C). Firstly, semithin sections were cut on Ultrotome Nova (LKB, Sweden) and stained with toluidine blue. Subsequently, ultrathin sections were cut on the same ultramicrotome, collected onto formvar carbon-coated copper grids, counterstained with uranyl acetate and lead citrate and examined under JEOL JEM-1400Plus transmission electron microscope (at 120 kV, JEOL, Japan).

### RNA

Two methods were employed to isolate RNA (i) Cells were collected for RNA isolation from 2D controls by washing the cells once with DPBS^−/−^, followed by scraping the cells into DPBS^−/−^. The resulting cell suspension was pelleted by centrifugation at 300 × g for 1 minute at room temperature. The supernatant was carefully removed and Trizol (Life Technologies) was added to lyse the cells. For organoids, RNA isolation was performed by removing 2 ml of suspension culture medium from the Erlenmeyer flasks and collecting the organoids by centrifugation at 300 × g for 5 minutes at room temperature. The supernatant was carefully removed and the cells were washed with 5 ml of DPBS^−/−^ and repelleted as previously. The DPBS^−/−^ was gently removed and Trizol added to lyse the cells. The Trizol samples were then either processed immediately for RNA isolation according to the manufacturer’s instructions or stored at -80℃ for subsequent processing. RNA was quantified using a NanoDrop ND-1000 Spectrophotometer (NanoDrop). (ii) For the vitamin K dependent enzyme analysis total RNA was isolated using the MagMAX™-96 Total RNA Isolation Kit on a MagMAX™ Express-96 Deep Well Magnetic Particle Processor as described by the manufacturer (both from Thermo Fisher Scientific, Waltham, MA, USA).

### cDNA synthesis

500 ng of RNA was used as a template for reverse transcription to cDNA. cDNA synthesis was performed using the High Capacity Reverse Transcriptase Kit (Life Technologies) with random primers, following the manufacturer’s instructions for reactions without RNase inhibitor.

### Gene expression analysis with RT-qPCR

Gene expression was analysed via reverse transcriptase quantitative polymerase chain reaction (RT-qPCR) using TaqMan probes (Life Technologies) and SSO Universal Probes Master Mix (Bio-Rad). For a complete list of probes used in this study, please see Supplementary Table 1. All samples were analysed in triplicate. Data is presented as the average of three independent experiments +/− the standard deviation.

### CYP450 activity and induction

Analysis of Cytochrome P450 (CYP) basal activity and inducibility was performed as previously described with several modifications for organoid cultures. Briefly the 2D HLCs and organoids were induced from day 20 onwards of differentiation with prototypical CYP450 inducers: for CYP3A4 we cultured with 25 μM Rifampicin and 100 μM Omeprazole for CYP1A2 (Sigma-Aldrich). Prior to starting the inductions the cells/organoids were washed with DPBS^−/−^ 4 times to remove hydrocortisone and dexamethasone. After washing the inducers were added to L-15 medium as described above, minus hydrocortisone and dexamethasone. The induction medium was refreshed every 24 hours for 3 days. 72 hours post induction the cells were assayed for CYP1A2 and CYP3A4 activity using the P450-Glo CYP3A4 (Luciferin-PFBE) Cell-based/Biochemical Assay kit (Promega, Cat. no. V8902) and the P450-Glo CYP1A2 Induction/Inhibition Assay kit (Promega, Cat. no. V8422) according to the manufacturer’s instructions. Data was normalised to 1 million hepatocytes and is presented as the average of three independent experiments +/− the standard deviation.

### Heroin metabolism

After 21 days differentiation, 50 organoids per well were loaded in triplicate into a 96-well plate and treated with culture media supplemented with 10 μM heroin for 1, 3, 6, and 24 hours. For controls, we used culture media without organoids; these were performed in parallel to measure heroin degradation throughout the experiment. To stop metabolism at each time point the samples were transferred to a new 96-well plate prefilled with formic acid (final conc. 0.1 M) along with internal standards. The samples were centrifuged for 10 minutes at 1000 × g at 4°C. The supernatants were transferred to auto-sampler vials and analysed for heroin, morphine, and M3G using an Acquity UPLC system (Waters, Milford, MA) coupled to a Xevo-TQS triple quadrupole mass spectrometer with an electrospray ionization interface (Waters) based on a method previously described ^54^. Data acquisition, peak integration, and quantification of samples were performed using MassLynx 4.0 SCN509 software (Waters Corp., Milford, MA, USA).

### F7 activity

FVII activity in the cell medium was determined using the Human FVII Chromogenic Activity Kit (Nordic BioSite AB, Täby, Sweden) according to manufacturer’s instructions. Primary hepatocytes (Thermo Fisher Scientific) were used as control.

### Western analysis of factor II

Intracellular levels of FII were determined by western blot (WB). Briefly hepatic organoids were lysed in T-PER™ buffer (Thermo Fisher Scientific) and lysates were collected by centrifugation at 8000 × g for 10 minutes. Equal amounts of proteins from lysates were separated by SDS-PAGE using Mini-PROTEAN^®^ TGX™ 10% Precast Gels (Bio-Rad, Hercules, CA, USA) before transfer onto a Sequi-Blot PVDF membrane (Bio-Rad) using the Mini Trans-Blot Electrophoretic Transfer Cell system (Bio-Rad). The membranes were incubated overnight at 4°C with the primary antibody anti-FII (Novus biologicals, Centennial, CO, USA) or Beta-actin (Sigma Aldrich, Saint Louis, MO, USA). The membranes were washed and then incubated with the appropriate horseradish peroxidase (HRP)-conjugated secondary antibody (Santa Cruz Biotechnology, Dallas, TX, USA) for 1 hour at room temperature. The blots were developed using Radiance Plus chemiluminescent substrate (Azure Biosystems, Dublin, CA, USA) and the signals quantified using the ImageQuant LAS-4000 mini Imager (GE Healthcare, Chicago, IL, USA). Primary hepatocytes were used as control.

### Serum protein analysis via ELISA

ELISAs for human Albumin (Bethyl Laboratories Inc cat# E80-129), human Alpha-1-anti-trypsin (Abcam cat# ab108799) and hepatocyte growth factor (Antibodies-online cat# ABIN624992) were used as described by the supplier. Colorimetric readings were taken at the specified wavelength on a SPECTRAmax PLUS 384. Values were derived from standards using appropriate lines of best fit generated on Softmax pro software. For clotting factors organoids were collected by centrifugation for 10 minutes at 300 × g and the cell medium was collected. Hepatic organoids were lysed in T-PER™ buffer (ThermoFisher) containing Halt protease and phosphatase inhibitor cocktail 1X (Thermo Fisher cat# 78440). FVII and FX antigen (FVIIAg and FXAg) were measured in the cell medium using FVII ELISA kit and FX ELISA kit (Abcam, cat# ab168545 and ab108832) respectively. Primary hepatocytes were used as control. The FVIIAg and FXAg levels (ng/ml) were normalized to 1×10^6^ cells and the ratio organoid/ primary hepatocytes was calculated.

### Oleic acid accumulation assay

1 M Oleic acid (Sigma Aldrich) was diluted with NaOH (Sigma Aldrich) and heated at 70°C for 30 minutes to form a 20 mM Sodium Oleate solution. This was then diluted with a 5% BSA/PBS solution at 37°C to form a 5 mM Sodium Oleate/BSA complex. This was further diluted to 300 μM in culture media and incubated with organoids for 5 days, changing the media each day. After the fatty acid treatment, the organoids were then incubated with 3.8 μM BODIPY 493/503 (Life Technologies) for 30 minutes in culture media at 37°C, then washed two times with PBS before replacing with fresh culture media containing DRAQ5 (Thermofisher) at 1:1500. Organoids were then imaged on a confocal as described above.

### Sample preparation, processing and data processing of proteomics data

#### Patient liver samples

Five patient samples were collected from liver explants from patients undergoing LTX at Oslo University Hospital. The samples were stored in liquid nitrogen. The regional ethics committee approved the use of the patient material (REK 2012-286) in accordance with the Declaration of Helsinki. All participants provided written informed consent.

Samples were processed as follows, the proteins were precipitated with acetone/TCA (Sigma Aldrich). The pellets were resuspended in 8 M Urea in 50 mM NH_4_HCO_3_, and the proteins were reduced, alkylated and digested into peptides with trypsin (Promega). The resulting peptides were desalted and concentrated before mass spectrometry by the STAGE-TIP method using a C18 resin disk (3M Empore). Each peptide mixture was analyzed by a nEASY-LC coupled to QExactive Plus (ThermoElectron, Bremen, Germany) with EASY Spray PepMap^®^RSLC column (C18, 2 μl, 100 Å, 75 μm × 25 cm) using a 120 minute LC separation gradient.

The resulting MS raw files were submitted to the MaxQuant software version 1.6.1.0 for protein identification. Carbamidomethyl (C) was set as a fixed modification and acetyl (protein N-term), carbamyl (N-term) and oxidation (M) were set as variable modifications. First search peptide tolerance of 20 ppm and main search error 4.5 ppm were used. Trypsin without proline restriction enzyme option was used, with two allowed mis-cleavages. The minimal unique+razor peptides number was set to 1, and the allowed FDR was 0.01 (1%) for peptide and protein identification. The Uniprot database with ‘human’ entries (October 2017) was used for the database searches.

Proteins with log2(intensity) > 10 average intensity value were defined as “expressed proteins”. Pearson correlation coefficient between liver and organoid was calculated from log2(intensity) with cor.test function in R. Differential expression of proteins between liver and organoid was defined with more than 2 fold change and p<0.05 by two-sided T test. Gene Ontology analysis was conduced with GOstats Bioconductor package ^55^. Multiple test correction was performed by Benjamin-Hochberg method with p.adjust function in R.

### Library preparation and data processing of scRNAseq

scRNA-seq libraries were prepared from the liver organoids at day 48 with Chromium Single Cell 3’ Reagent Kits (version 2 - 10x Genomics) as described previously ^56^. Conversion to fastq format, mapping/UMI counting in human genome (hg19) and data aggregation were implemented by *mkfastq*, *count* and *aggr* functions with default parameters in CellRanger software (v2.1.0). Subsequent data processing, such as batch effect normalization, was performed by Seurat software (v3.1.0)^57^. In each replicate, the feature UMI count was normalized to the total count and multiplied by 10,000. Top 2,000 Highly-Variable Features (HVFs) were then identified by variance stabilizing transformation. Anchor cells across different scRNAseq libraries were identified with HVFs under 20 dimensional spaces from canonical correlation analysis and used for the transformation of multiple scRNAseq datasets into a shared space. Gene expression values were scaled for each gene across all integrated cells and used for principal component analysis (PCA). 20 PCs were further assigned into two dimensional space using Uniform Manifold Approximation and Projection (UMAP) and also used to identify cell clusters. Differentially-expressed genes in each cluster were identified with more than 1.25 fold change and p<0.05 by two-sided T test. Overrepresented GO terms were identified by GOstats (v2.24.0)^55^. Multiple test correction was performed by Benjamin-Hochberg method with p.adjust function in R.

The cluster labels were assigned by cell type specific markers and GO terms (Supplementary Fig. 2c,e). Nine out of 22 clusters were first separated by the overrepresentation of “extracellular matrix (GO:0031012)”, which is a feature of stellate and endothelial cells. Active (AST) and resting stellate cells (RST) were defined by genes involved in “mitotic nuclear division (GO:0140014)” and its markers (MGP and ELN). Non-stellate clusters were labeled as endothelial cell (EC) and further divided into liver sinusoidal (LSEC) and macrovacular endothelial cells (MVEC) by the absence and presence of vasculogenesis markers (KDR and HAND1)^58^. Seven out of 13 other clusters were assigned hepatocyte (HEP), cholangiocyte (CHO), Kupffer cell (KPC) and Kupffer precursors (KPP) using the enrichment of GO terms “Cholesterol homeostasis (GO:0042632)”, “Keratinization (GO:0031424)”, “Phagocytosis (GO:0006909)” and “hematopoietic stem cell differentiation (GO:0060218)”, respectively. Five clusters were assigned as peripheral nervous system with the expression of neuronal lineage markers (SOX2 and PAX6) and further divided into neuron (Neu), glia (Glia), neuro progenitor (NPC) and cilia-bearing cell (CBC) with “axon development (GO:0061564)”, “glial cell development (GO:0010001)”, “mitotic nuclear division (GO:0140014)” and “cilium assembly (GO:0060271)”, respectively ^56^. We could not identify any unique marker and relevant GO terms in one cluster and labeled it as unknown (UN). The cluster labelling strategy was schematically represented in Supplementary Fig. 2b.

Public transcriptome profiles were downloaded from NCBI Gene Expression Omnibus database. Single-cell transcriptome of the liver organoid from Ouchi *et al*. (GSE130073)^24^ and human liver atlas (GSE124395)^25^ were merged with our scRNAseq data and plotted into the share UMAP space by Seurat as described above. Clusters, which are mainly composed of CD45^+^ cells and unique to human liver atlas, were labeled as “other immune cell”. Genes were sorted by the difference of average expression of all cells between our and Ouchi et al. liver organoid and used for GSEA (v2.2.2) of REACTOME genes without collapsing gene set ^59^. Cell-type specific gene signatures were constructed from bulk RNA-seq in primary hepatocyte (GSE98710, GSE112330 and GSE135619)^11, 12^, biliary tree stem (GSE73114)^13^, stellate (GSE119606)^14^ and endothelial cells (GSE114607)^15^. The RNA-seq read was aligned to hg19 human genome by Tophat (v2.2.1) with default parameters^60^. The mapped reads were counted in each gene by HTSeq software (v0.9.0) with options “-s no -f bam”^61^.

The factors of technical variations across multiple transcriptome datasets were minimized by RUVs function in RUVSeq (v1.8.0)^62^. Subsequently, differentially expressed genes in each cell type were identified by DESeq2 (v1.14.1). To evaluate the enrichment of the cell-type specific genes, genes were sorted in individual cells by relative expression level to average of all cells and used for GSEAPY software (v0.9.3) with options “--max-size 50000 --min-size 0 -n 1000”. Hepatic zone-specific genes were obtained from transcriptome profiles of hepatocyte from laser-microdissected human livers (GSE105127)^18^. After processing the bulk RNA-seq, the zone-specific genes were defined with more than 1.5-fold change and p<0.05 by two-sided T test. The enrichment was evaluated by GSEAPY software with pre-ranked genes in individual cells relative to all cells in all hepatocyte clusters.

To investigate transcriptional bias between LSEC and MVEC, cells from EC clusters were ordered in pseudotemporal spaces by Monocle (v2.99.3). Briefly, the monocle object was first constructed from the UMI count matrix for cells in EC clusters and preprocessed according to the instruction. We then replaced data in “normalized_data_projection” and “reducedDimW” with non-transposed and transposed PCA dimensional matrix. In addition, “reducedDimS”, “reducedDimA” and “reducedDimK” slot were replaced with transposed UMAP dimensional matrix. The principal graph was learned by learnGraph function with “RGE_method =’DDRTree’, close_loop=T, prune_graph=F, euclidean_distance_ratio=5”. Subsequently, cells are ordered according to the trajectory by orderCells function using MVEC1 as a root cluster. Differentially-expressed genes were identified by differential GeneTest function with the model “~sm.ns(Pseudotime)”. Finally. Genes with q<1e-50 were selected as EC ordering-dependent genes and used for GO analysis by GOstats as described above.

### Animal work

Male and female NOD.Cg-Prkdc^scid^ Il2rg^tm1Wjl^/SzJ (NOD *scid* gamma, NSG) mice (purchased from The Jackson Laboratory, Bar Harbor, ME, USA) were housed in a Minimal Disease Unit at the animal facility at Oslo University Hospital Rikshospitalet, Oslo, Norway, with a 12 hour light–dark cycle and *ad libitum* access to water and standard rodent diet. All experiments were performed with co-housed age-matched mice. Mice undergoing surgery were not fasted and were 15 weeks of age at the time of surgery. All animals received human care and the animal experiments were approved the Norwegian National Animal Research Authority (project license no FOTS 19470) and performed according to the European Directive 2010/63/EU, the Animal Research: Reporting of In Vivo Experiments guidelines and The Guide for the Care and Use of Laboratory Animals, 8th edition (NRC 2011, National Academic Press).

### Implantation of human liver organoids under the rodent kidney capsule

Male and female immunodeficient NSG mice were used in this study. The transplantation of the organoids under the kidney capsule or the sham laparotomy was performed as described in ^63^ with some modifications. In brief, the procedure was performed using proper multimodal analgesia with s.c. administration of an local analgesia (Marcain, 0.07 ml/10 g BW) in combination with s.c. administration of Buprenorfin (0.1 mg/kg) before surgery and general anesthesia with i.p. injection with FD2 (Fentanyl/Domitor/Dormicum) and Antisedan (antagonist) post surgery. Following a sterile preparation of the left flank, a 1.5 cm incision was made midway between the last rib and the iliac crest and approximately 0.5 cm parallel and ventral to the spine ^63, 64^. The left kidney was slowly externalized through the abdominal incision using sterile cotton swabs, immobilized using non-traumatic forceps and moisturized with warm sterile saline. The injection site was located at the upper lateral side of the kidney, and a 1 ml syringe with a 25G needle containing either the organoid suspension or pure Matrigel (Thermo Fisher Scientific) (sham surgery) was gently pushed under the capsule towards the inferior pole of the kidney in order to avoid perforation and damage to the blood vessels. 50-80 μl organoid suspension or Matrigel matrix was very slowly discharged under the kidney capsule and the needle simultaneously slowly pulled out of the capsule to avoid backflow. Next, the kidney was returned to the body cavity, the abdominal wall closed with suture and the skin incision closed with 7 mm wound clips. Post surgery, the mice were examined daily the first week for normal wound healing, with weight measurements and general well-being, thereafter once a week. Once every week, blood was sampled from the saphena vein for serum markers measurement.

## Supplementary Materials

Fig. S1. Organisation of D7 organoids.

Fig. S2. Cell Type Characterization in Single-Cell Transcriptome Profiles.

Fig. S3. Organisation and Secretion in D20-D30 Organoids.

Fig. S4. A Scaleable Culture System.

Table S1. List of Reagents

## Acknowledgments

We would like to thank the confocal microscopy core facility Rikshospitalet, Gaustad node, Oslo for use of their equipment.

## Author contributions

GJS, SPH, RS and IHP designed and executed the development of the study. GJS, TY, IHP, YX and BP prepared libraries, ran single cell RNA seq. TY, IHP, FSS, TAN and GJS performed single cell RNA sequencing bioinformatical analysis and proteomic analysis. Proteomic data acquisition was performed by TAN, SRW, RA and FSS. Establishment of suspension system SPH, RS, GJS, RZ, and HK. Metabolism data acquisition RS, SPH, SRW, FSS, ICB and ILB. Serum protein and coagulation factor data acquisition GJS, SPH, RS, MEC, PMS and EA. Modeling organoid growth JJW. Electron microscopy and immunohistochemistry data acquisition GJS, SPH, AB, DC and JM. Transplantation work and data acquisition EM, KSÅ, EA and SPH. Other data acquisition was performed by SPH, RS, SFB, SM, KS, FB, DK, RA. GJS, SPH and RS wrote the manuscript. All authors critically evaluated the manuscript.

