## Supplemental data and table for "Scalable production of tissue-like vascularised liver organoids from human PSCs"

Supplementary Materials:

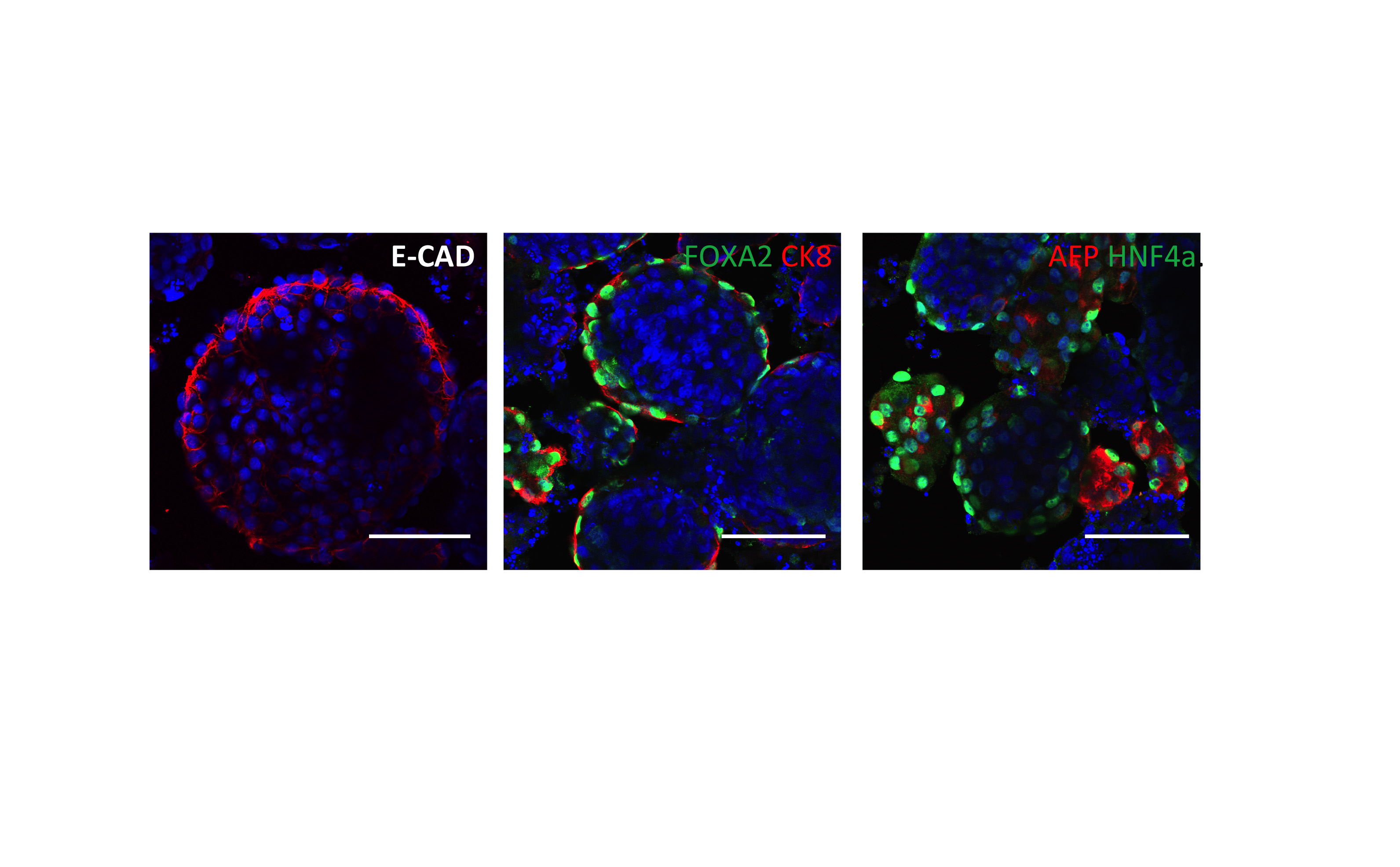

**Supplementary Figure 1**. Organisation of D7 organoids. Whole-mount immunostaining showing a cross section through the D7 organoids showing the restriction of epithelial (ECAD) and early hepatocyte markers (FOXA2, CK8, HNF4, AFP) to the outer surface of the organoids and their absence from the inner core. Nuclei stained with DRAQ5, scale bar = 100 µm.

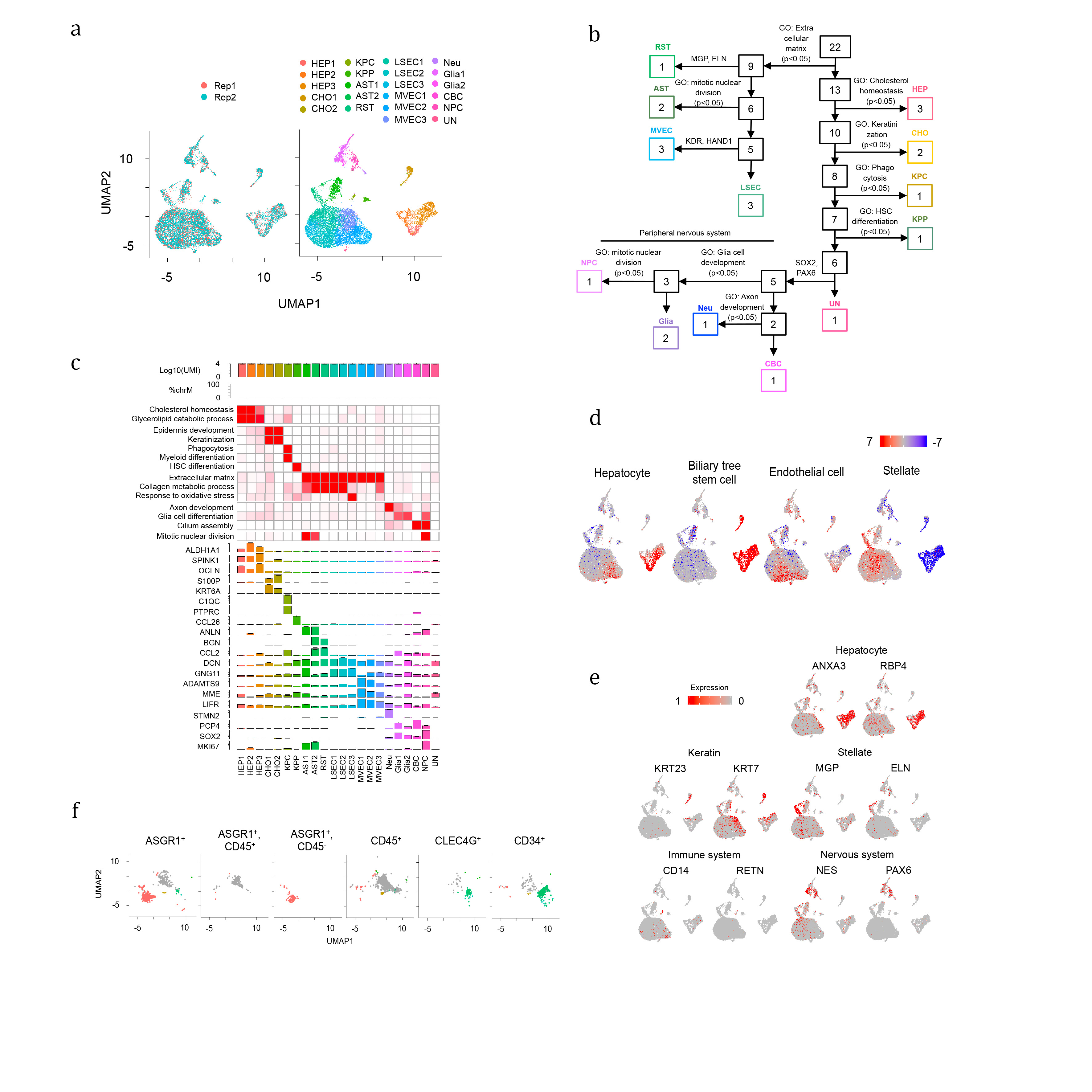

**Supplementary Figure 2**. Cell Type Characterization in Single-Cell Transcriptome Profiles. (A) UMAP plot of single cells distinguished by replicates and clusters. (B) Schematic representation of cluster annotation method. (C) Comparison of total UMI, percentage of mitochondria-derived reads, GO enrichment, representative gene markers across 22 clusters. (D) GSEA of gene signatures of hepatocyte, biliary tree stem cell, endothelial cell and stellate. The enrichment and depletion are scaled by -log10(FDR) and shown by red and blue colors, respectively. (E) Expression pattern of cell-type specific markers in the liver organoid. Relative expression level is plotted from gray to red colors. (F) UMAP plots of FACS-sorted cells from human livers. Color scheme and UMAP dimension correspond to Fig. 2E.

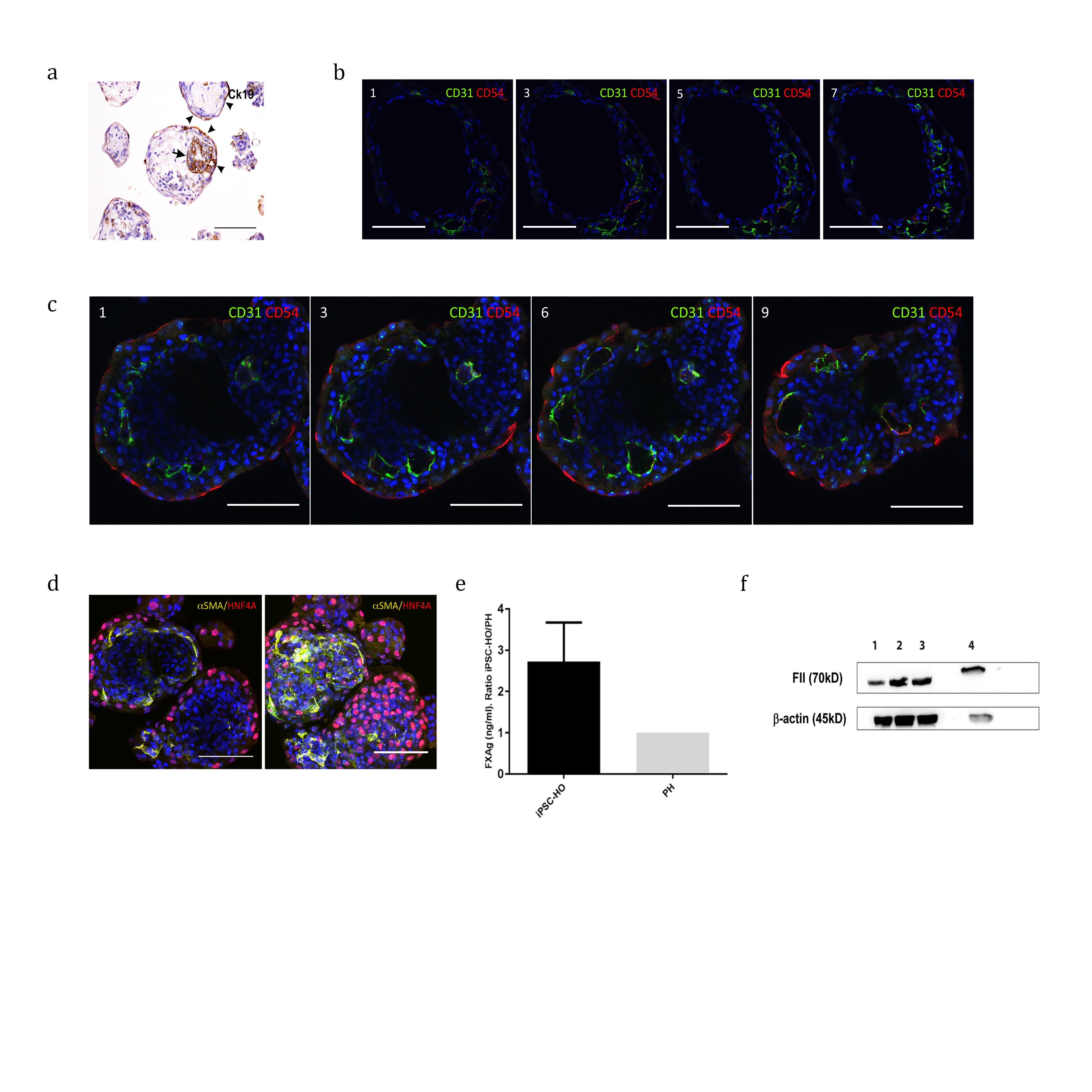
**Supplementary Figure 3**. Organisation and Secretion in D20-D30 Organoids. (A) Immunohistochemical staining of CK19 positive cells, arrow denotes cholangiocytes surrounding lumen. (B) and (C) z planes covering a volume of 35 μm (B) or 45 μm (C) from cryosections immunostained (same organoid as Fig. 5A) for the endothelial markers CD31 and CD54 showing continuity of luminal spaces and changes in endothelial marker expression towards the outer surface of the organoids. (D) Whole-mount immunostaining showing SMA positive cells beneath the hepatocyte layer. (E) FX protein (Ag) levels (ng/ml) in culture medium from iPSC-HO and PH as measured by ELISA. The total concentration of FX was adjusted to 1x10^6^ cells and the results are expressed as the ratio iPSC-HO/PH. Results from three independent experiments are presented as mean ±SEM. (F) Lysates were obtained from iPSC-HO (lanes 1 -3) and PH (lane 4). Equal amounts of proteins were separated by SDS-PAGE under reducing conditions, blotted onto a PDFV membrane and incubated with antibody against FII. β-actin was used as loading control. Results of three independent experiments are presented. All Scale bars are 100 µm unless stated otherwise.

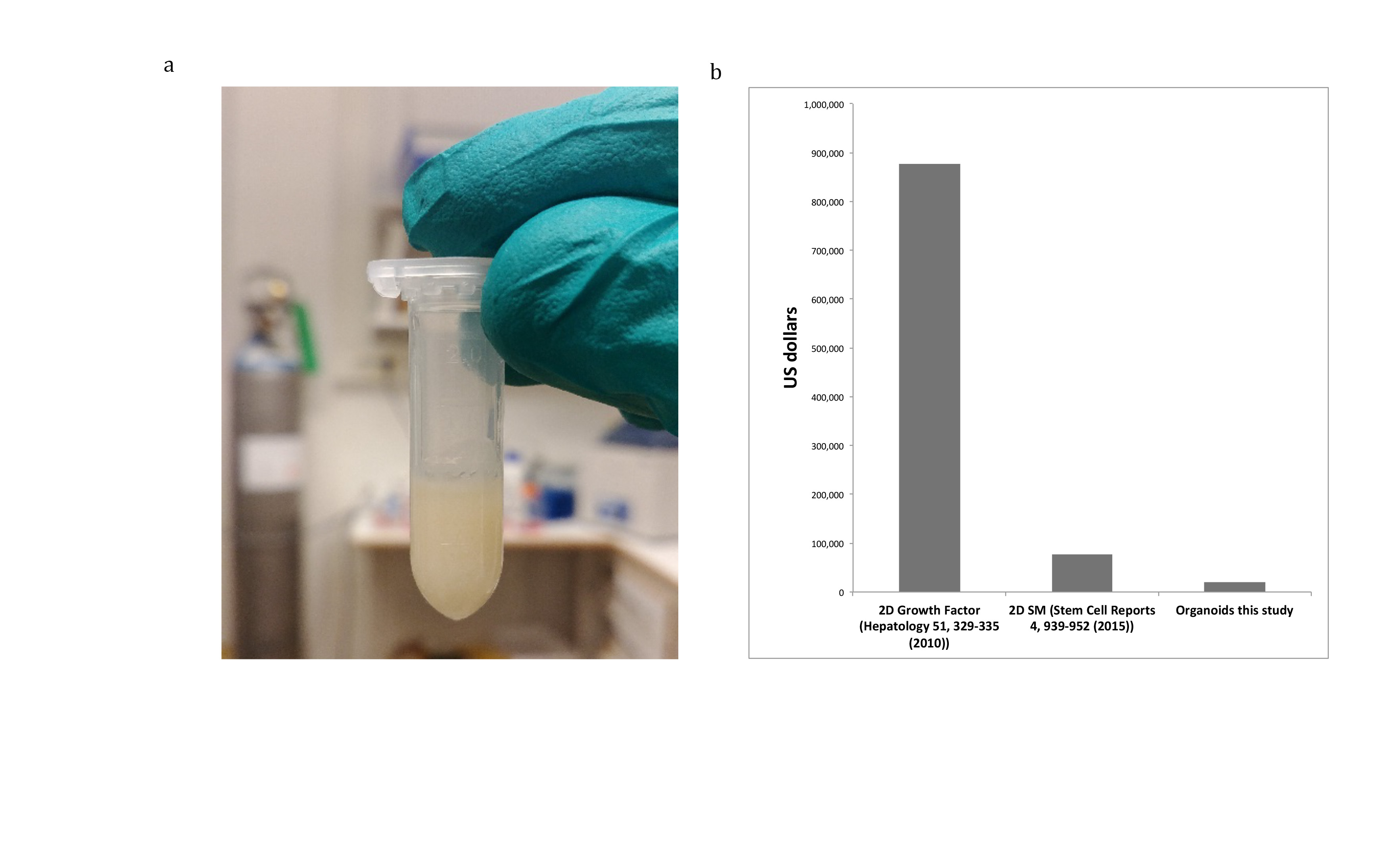
**Supplementary Figure 4**. A Scaleable Culture System**.** (A) Image showing pellet of end stage organoids from 35 ml of suspension culture after centrifugation, approximately 18,000 organoids in a 700 µl packed pellet. (B) Cost comparison in US dollars for the production of approximately 2x10^10^ cells based on a previously published methods using growth factor in 2D ^5^, a small molecule-based differentiation ^7^ and this study.

**Supplementary Table 1**

| **Taqman assay** | **Catalog number** | **Supplier** |
| --- | --- | --- |
| Hs01551992_m1 (F7) | 4331182 | Thermo Fisher Scientific |
| Hs01011988_m1 (F2) | 4331182 | Thermo Fisher Scientific |
| Hs00252034_m1 (F8) | 4331182 | Thermo Fisher Scientific |
| Hs01592597_m1(F9) | 4331182 | Thermo Fisher Scientific |
| Hs00173450_m1 (F10) | 4331182 | Thermo Fisher Scientific |
| Hs00165584_m1 (PROC) | 4331182 | Thermo Fisher Scientific |
| Hs00170586_m1 (FGB) | 4331182 | Thermo Fisher Scientific |
| Hs00166654_m1 SERPINC1 (antithrombin) | 4331182 | Thermo Fisher Scientific |
| Hs01097800_m1 SERPINA1 (alpha 1 antitripsin) | 4331182 | Thermo Fisher Scientific |
| Hs00230853_m1 (HNF4A) | 4331182 | Thermo Fisher Scientific |
| Hs99999901_s1 (18S) | 4331182 | Thermo Fisher Scientific |
| Hs01005019_m1 (ASGR1) | 4331182 | Thermo Fisher Scientific |
| Hs01053049_s1 (SOX2) | 444892 | Thermo Fisher Scientific |
| Hs04260366_g1 (NANOG) | 444892 | Thermo Fisher Scientific |
| Hs00430824_g1 (MIXL1) | 444892 | Thermo Fisher Scientific |
| Hs00232764_m1 (FOXA2) | 444892 | Thermo Fisher Scientific |
| Hs00751752_s1 (SOX17) | 444892 | Thermo Fisher Scientific |
| Hs00242160_m1 (HHEX) | 444892 | Thermo Fisher Scientific |
| Hs00173490_m1 (AFP) | 444892 | Thermo Fisher Scientific |
| Hs00269972_s1 (CEBPA) | 444892 | Thermo Fisher Scientific |
| Hs00171403_m1 (GATA4) | 444892 | Thermo Fisher Scientific |
| Hs00426361_m1 (CYP3A7) | 444892 | Thermo Fisher Scientific |
| Hs00910225_m1 (ALB) | 444892 | Thermo Fisher Scientific |
| Hs00199611_m1 (TDO2) | 444892 | Thermo Fisher Scientific |
| Hs00610080_m1 (T) | 4331182 | Thermo Fisher Scientific |
| Hs00896293_m1 (PROX1) | 4331182 | Thermo Fisher Scientific |
| Hs00195612_m1 (TBX3) | 4331182 | Thermo Fisher Scientific |
| Hs00999634_gH (POU5F1) | 4331182 | Thermo Fisher Scientific |
| Hs00906630_g1 (GSC) | 4331182 | Thermo Fisher Scientific |
| Hs00415443_m1 (NODAL) | 4331182 | Thermo Fisher Scientific |
| Hs00193796_m1 (CER1) | 4331182 | Thermo Fisher Scientific |
| Hs00846731_s1 (SOX7) | 4331182 | Thermo Fisher Scientific |
| Hs00174914_m1 (TTR) | 4331182 | Thermo Fisher Scientific |
| Hs00952079_g1 (APOA2) | 4331182 | Thermo Fisher Scientific |
| Hs00604506_m1 (CYP3A4) | 4331182 | Thermo Fisher Scientific |

Primary Antibodies, Secondary Antibodies and Serum

| **Marker** | **Supplier** | **Catalogue No.** | **Species** | **Dilution** |
| --- | --- | --- | --- | --- |
| αSMA | Invitrogen | MA5-11547 | Rabbit | 1:400 |
| ALB | Sigma Aldrich | A6684 | Mouse | 1:600 |
| ALCAM | Santa Cruz | sc-74558 | Mouse | 1:200 |
| AFP | Sigma Aldrich | A8452 | Mouse | 1:600 |
| ASGPR1 | Santa Cruz | sc-5623 | Mouse | 1:200 |
| B3 Tubulin | Santa Cruz | sc-80005 | Mouse | 1:200 |
| Beta-actin | Sigma Aldrich | A5441 | Mouse | 1:5000 |
| CD31 | Abcam | ab32457 | Rabbit | 1:1000 |
| CD34 | Santa Cruz | sc-7324 | Mouse | 1:200 |
| CD54 | Thermofisher | MA5-1301 | Mouse | 1:100 |
| CD68 | Santa Cruz | sc-20060 | Mouse | 1:300 |
| CK19 | Amersham | RPM1165 | Mouse | 1:150 |
| CK7 | DAKO | M7018 | Mouse | 1:100 |
| CK8 | DAKO | M0631 | Mouse | 1:100 |
| CPS1 | Santa Cruz | sc-376190 | Mouse | 1:150 |
| CYP2A6 | Origene | TA503832 | Mouse | 1:100 |
| ECAD | Santa Cruz | sc-2179 | Mouse | 1:75 |
| FII | Novus Biologicals | NBP1-58268 | Rabbit | 3µg/ml |
| FOXA2 | Abcam | Ab108422 | Rabbit | 1:300 |
| GS | Transduction Laboratories | G45020/L1 | Mouse | 1:250 |
| HNF4α | Santa Cruz | sc-8987 | Rabbit | 1:300 |
| LAM | Thermofisher | PA5-16287 | Rabbit | 1:250 |
| LYVE1 | Abcam | ab33682 | Rabbit | 1:200 |
| MESP1 | Thermofisher | PA5-67086 | Rabbit | 1:300 |
| MRP2 | Abcam | ab3373 | Mouse | 1:100 |
| Nanog | Stemgent | 09-0020 | Rabbit | 1:100 |
| OCT4 | Stemgent | 09-0023 | Rabbit | 1:100 |
| SOX2 | Stemgent | 09-0024 | Rabbit | 1:100 |
| WT1 | Thermofisher | PA5-16879 | Rabbit | 1:300 |
| ZO1 | Santa Cruz | sc-33725 | Rat | 1:50 |
| Alexaflour'488'anti-rabbit | Life Technologies | A21206 | Donkey | 1:1500 |
| Alexafluor'488'anti-mouse | Life Technologies | A11059 | Rabbit | 1:1500 |
| Alexafluor'594'anti-mouse | Life Technologies | A11005 | Goat | 1:1500 |
| Alexafluor'488'anti-rat | Life Technologies | A11006 | Goat | 1:1500 |
| Mouse anti rabbit (HRP) | Santa Cruz | sc-2357 |  | 1:10000 |
| m-IgGκ BP-HRP | Santa Cruz | sc-516102 |  | 1:10000 |
| Goat serum | Sigma | G9023 |  |  |
